# Integration of sleep homeostasis and navigation in *Drosophila*

**DOI:** 10.1101/2020.07.23.217638

**Authors:** Andres Flores Valle, Pedro J. Gonçalves, Johannes D. Seelig

## Abstract

During sleep, the brain undergoes dynamic and structural changes. In *Drosophila*, such changes have been observed in the central complex, a brain area important for sleep control and navigation. The connectivity of the central complex raises the question about how navigation, and specifically the head direction system, can operate in the face of sleep related plasticity.

To address this question, we develop a model that integrates sleep homeostasis and head direction. We show that by introducing plasticity, the head direction system can function in a stable way by balancing plasticity in connected circuits that encode sleep pressure. With increasing sleep pressure, the head direction system nevertheless becomes unstable and a sleep phase with a different plasticity mechanism is introduced to reset network connectivity.

The proposed integration of sleep homeostasis and head direction circuits captures features of their neural dynamics observed in flies and mice.

## 1 Introduction

Sleep affects many different brain functions such as cognition^1^ or working memory^2^ and sleep dysfunction is related to a range of diseases^3^. Sleep is observed across species and different hypotheses have been put forward to explain the function of sleep^4^, for example reverse learning of spurious network states (that were created as a byproduct of intended memories)^5-7^ or weakening of synapses (synaptic homeostasis hypothesis)^8^.

The function of sleep is linked to the dynamic and structural changes that it induces in the brain^9^, which in turn are monitored by sleep control or sleep homeostasis circuits^10,11^. The circuits that control sleep are distributed over many different brain areas and cell types^10^. Thanks to the genetic tools^12,13^ that allow dissecting neural circuits into small populations of genetically identified cell types, as well as more recently the fly connectome^14^, *Drosophila* has emerged as a valuable model for sleep control^11,15-17^.

A generic sleep control circuit has been linked to specific neural populations in the brain of *Drosophila* in^18^. This circuit has three components and corresponding neural populations in the central complex (Figure 1A). A first component encodes sleep pressure. The corresponding neural population has been identified in the so called R5 ring neurons which arborize in concentric rings in the ellipsoid body^19^, a substructure of the central complex. These R5 neurons increase both their activity and synaptic strength over waking time and are reset with sleep. A second component of the sleep control circuit executes the switch between sleep and wakefulness (depending on the amount of sleep pressure). The corresponding neural population has been associated with the the dorsal fan-shaped body (dFB) neurons, which promote sleep when active^20^. A third component triggers locomotion, processes visual input, and increases sleep pressure^18^ and the corresponding neurons are so called helicon cells^18^, also identified as ExR1 neurons^21^. The proposed recurrent circuit between these three neural populations^18^ is illustrated in Figure 1A.

**Figure 1.**
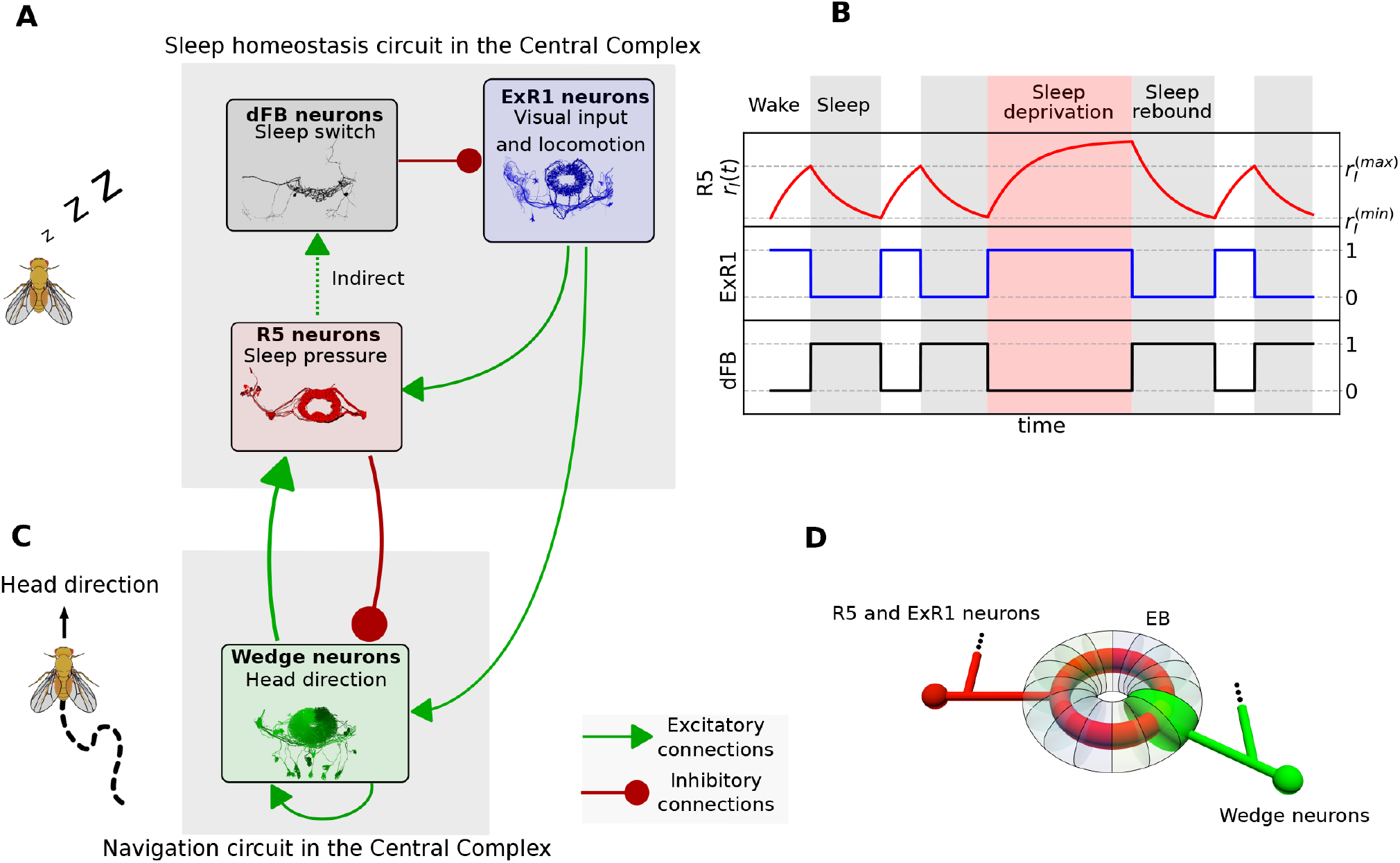
Sleep homeostasis and navigation circuits in the central complex. **A** Recurrent sleep homeostasis circuit proposed in^18^. The three populations are connected via excitatory (green arrows) or inhibitory (red arrows) connections. **B** Simulation of the sleep homeostasis model illustrating the dynamics of each population over time. **C** Interaction between the fly head direction circuit and populations involved in sleep homeostasis. **D** Schematic of connectivity between wedge neurons and R5 and ExR1 neurons in the ellipsoid body. Images are downloaded from the connectome database^14^.

The same central complex structures involved in sleep have also been shown to be important for navigation. In particular, ring neurons with similar morphology to the sleep-related R5 ring neurons, provide sensory input to the head direction system, such as visual features^22,23^ or wind direction^24^. Such input is integrated in so called wedge neurons, which arborize in different wedges of the ellipsoid body, where they intersect with ring neurons. These wedge neurons encode the head direction of the fly through a bump of activity that moves around the ellipsoid body^25^.

In the context of navigation, the structure and function of circuits in the central complex are reminiscent of ring attractor networks^25^. Such networks, which are well suited to encode a circular variable, have been suggested to underlie the encoding of head direction, originally in mammals^26^ and more recently in flies^25^.

It is currently unknown why the circuits for sleep homeostasis and head direction converge in the central complex. The morphological similarity of the ring neurons involved in sleep and head direction and the spatial proximity of the circuits as well as the fly connectome^14^, suggest that they interact. Given the observed activity and structural changes in R5 ring neurons after prolonged waking time and after sleep^19,27,28^, this suggests that the head direction system in the ellipsoid body needs to operate in the face of substantial synaptic and functional changes in connected circuits.

Motivated by this interaction between navigation and sleep homeostasis circuits as well as their plasticity^19,27,28^, we here use theoretical modeling to investigate how these two circuits can be understood as a combined system. For this purpose, we first model the circuit proposed in^18^ and confirm that it generates sleep homeostasis. We then extend the model by combining it with a head direction network as suggested by the connectome. In this combined model, the sleep pressure encoding R5 neurons balance Hebbian plasticity introduced in the recurrent connections of the head direction system. In this way, ring neurons maintain a functioning head direction system and record sleep pressure. The system is finally reset through a sleep phase.

We discuss how this model can integrate several experimental observations on the navigation and sleep homeostasis systems reported in the literature. We further discuss several predictions of the model that can be tested in experiments. This analysis contributes to an understanding of the generation and dynamics of sleep drive and links the control of sleep to sleep function.

## 2 Results

### 2.1 Sleep homeostasis model

The sleep homeostasis model proposed in^18^ is illustrated in Figure 1A. All connections between populations are direct^18^, except the connection between R5 and dFB neurons, which is considered indirect since these neural populations are not anatomically, but functionally connected^19^. This circuit is described by the following phenomenological model:

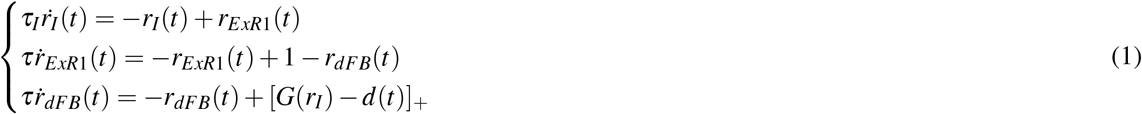

The variables *r_I_*(*t*), *r_ExR1_*(*t*) and *r_dFB_*(*t*) represent population firing rates of R5 neurons, ExR1 neurons and dFB neurons, respectively. The time constant *τ_I_* accounts for the slow dynamics of increasing activity observed during waking time in R5 neurons^19^ on the order of hours. The effective population time constant *τ* accounts for neural dynamics in the millisecond range. [·]_+_ is a threshold-linear function to ensure positive-valued firing rates. For simplicity, the model is defined such that population firing rates are between 0 and 1. The variable *d* (*t*), which can take values 0 or 1, represents an input to dFB neurons such as a wake-promoting dopaminergic signal^29^. The function *G*(*r_I_*), which depends on the history of activity of R5 neurons and produces the observed switching behavior in dFB neurons^18^, is described with a simple hysteresis (equation (9)).

Figure 1B shows a simulation of this model with the firing rates of the different populations changing over time. The sleep and wake phases are defined in terms of the activity of dFB neurons, which promote sleep while active^20^. During the wake phase, activity in R5 neurons increases, encoding sleep pressure due to sustained constant input from ExR1 neurons. After ring neurons reach an upper threshold, 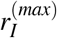, dFB neurons ‘switch on’ and inhibit activity in ExR1 neurons, which leads to a decrease in activity of R5 neurons. Once R5 activity reaches a lower threshold, 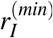, dFB neurons switch off, repeating the cycle. Sleep deprivation (by setting *d*(*t*) = 1 in the top red region) inhibits activity of dFB neurons and activity of ring neurons increases beyond 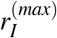. As expected for a sleep homeostasis circuit, after sleep deprivation, more time is required to reset the activity of R5 neurons back to 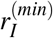 (sleep rebound, see Methods 4.3 for details of simulation).

### 2.2 Connectivity between head direction and sleep circuits

While the circuit described above can produce sleep homeostasis, the connectome^14^ shows that it acts not in isolation but interacts with the head direction system. Figure 1A and C show how R5 and ExR1 neurons are connected to wedge neurons that encode head direction. The anatomical organization of wedge, R5 and ExR1 neurons is shown schematically in Figure 1D, where each wedge neuron arborizes in a different wedge along the ellipsoid body, and R5 and ExR1 neurons arborize in concentric rings. The wedge neurons that encode head direction have been identified as EPG neurons^25^, but a similar population of wedge neurons, called EL^14, 30^ or AMPG-E^31^, could also potentially encode head direction. These neurons have been proposed to contribute to the persistent activity in the network by excitatory feedback to EPG neurons^31^. These neurons can mediate a connection between R5 and EPG neurons that is stronger than the direct connection between R5 and EPGs. In the following, wedge neurons refer to both EPG or EL populations without distinction. We assume that both encode head direction and are directly or indirectly connected to R5 neurons. In Figure S1, we show recurrent connections between wedge neurons (Figure S1A), between wedge and R5 neurons (Figure S1B)), and between wedge and ExR1 neurons (Figure S1C) according to the connectome^14^.

### 2.3 Integration of sleep homeostasis and ring attractor circuits without plasticity

The interaction of the sleep homeostasis and ring attractor circuits extracted from the fly connectome is shown schematically in Figure 2 A. Given that R5 neurons and wedge neurons are bidirectionally connected (see Figure S1B), we first asked how increasing activity of R5 neurons during the wake phase (see Figure 1B, first row) affects the head direction network. We therefore combined a ring attractor network with the above sleep homeostasis model (section 2.1) according to the connectivity in Fig. 2 A. As in previous work^32, 33^, we identify wedge neurons as the excitatory component of a ring attractor network with recurrent excitation, encoding head direction with sustained bump-like activity. On the other hand, we assume that R5 neurons provide increasing inhibition to wedge neurons^34^, in agreement with the majority of ring neurons being inhibitory^35,36^. (For simplicity, we assume that ExR1 neurons, which are bidirectionally connected to wedge neurons, provide input to the ring attractor similar to other ring neurons, and that wedge neurons and not ExR1 neurons, as suggested in^18^, charge the sleep homeostat.)

**Figure 2.**
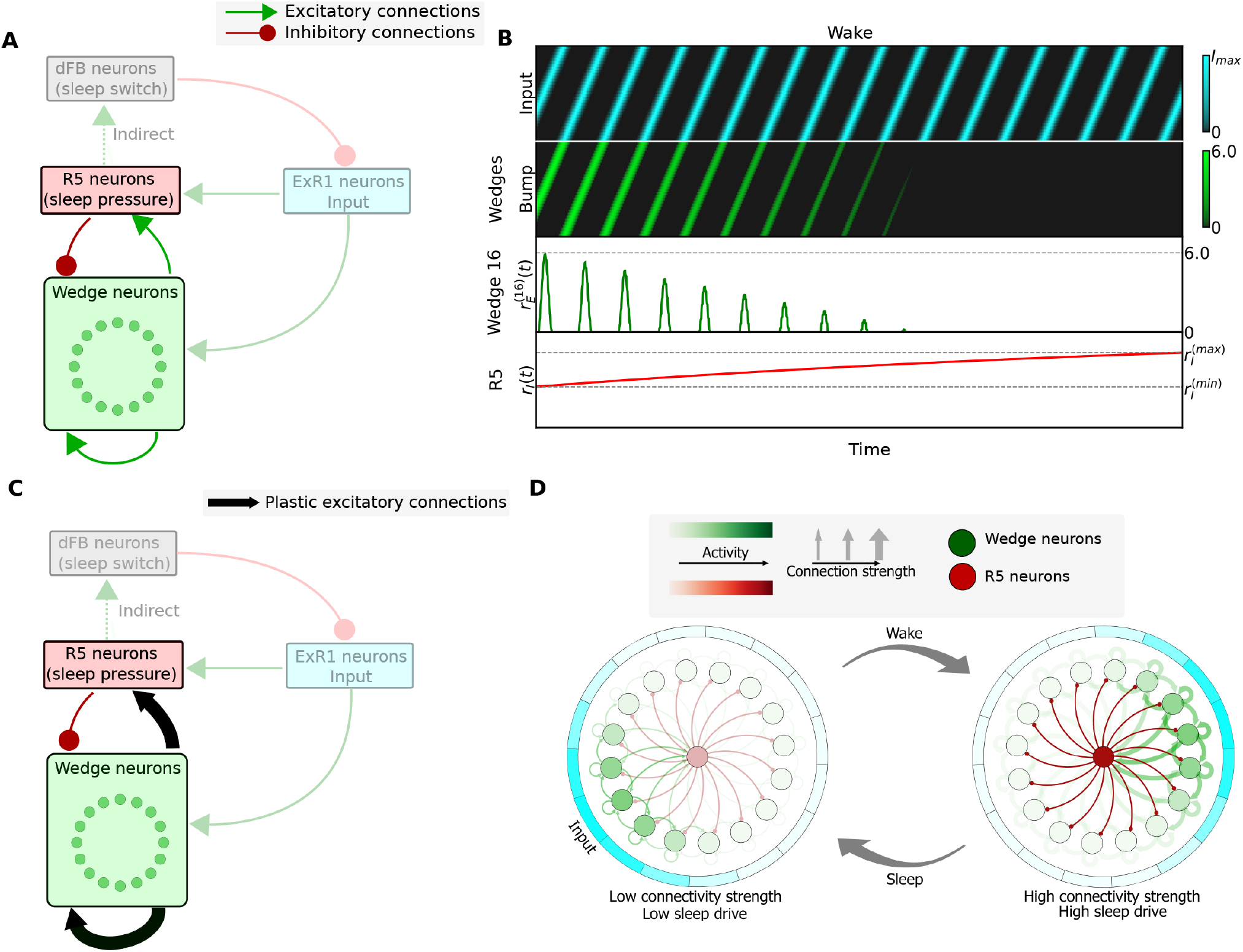
Integration of the sleep homeostasis circuit with a ring attractor network. **A** Schematic of a model where wedge neurons are connected to the sleep homeostasis circuit according to the fly connectome (see Figure S1). In this model, fixed connections are assumed. **B** Simulation of the model in A (dFB neurons not shown). Top row: rotating input to the ring attractor with a frequency 0.5 Hz. Second row: bump of activity in wedge neurons encoding head direction. Third row: activity of the wedge neuron 16 (representative for any other wedge neuron). fourth row: increasing activity of R5 neurons. **C** Model with plastic connections indicated by black arrows. **D** Dynamics of the model with plasticity: after a wake phase, high connectivity strength in the ring attractor leads to high sleep drive in R5 neurons, which leads to a switch to the sleep phase. After sleep, connectivity strength in the ring attractor is reset, producing low sleep drive.

Figure 2B shows the activity of wedge neurons and R5 neurons with a rotating input, representing visual or idiothetic cues, (in blue, first row) which as expected moves the bump around the ring attractor (in green, second row). Increased activity in R5 neurons, as experimentally observed with increased sleep drive, decreases the bump amplitude in the ring attractor until it finally vanishes. Therefore, simply connecting R5 and wedge neurons as indicated by the connectome, leads to a decreasing bump of activity over time.

### 2.4 Integration of sleep homeostasis and ring attractor circuits with plasticity

In Figure 2C, we propose an alternative model that can sustain a stable bump amplitude. In order to overcome a decreasing bump amplitude (which has not been experimentally observed), we hypothesize that the increase of inhibition from R5 neurons, in addition to encoding sleep drive, has the role of compensating for an increase in excitation in the head direction circuit. In particular, we hypothesize that excitatory synaptic strength between wedge neurons increases during the wake phase. This could be due to Hebbian plasticity between wedge neurons, since encoding the head direction in a bump of activity requires several wedge neurons to be active at the same time, thus strengthening the recurrent connections. This model is additionally motivated by the experimentally observed increase of activity as well as plasticity in R5 neurons^19^. In agreement with these data, we additionally add Hebbian plasticity from wedge neurons to R5 neurons.

In this model, R5 neurons act as a closed-loop feedback controller that prevents activity in wedge neurons from increasing due to hebbian plasticity, by adaptively increasing inhibition. In the context of the sleep homeostasis circuit, dFB neurons then detect synaptic growth in R5 neurons. When R5 neural activity reaches an upper threshold, dFB neurons switch on sleep. We assume that during sleep, the plastic connections are reset with LTD, decreasing activity in R5 neurons (as observed in^19^) and lowering sleep drive (Figure 2D, left).

The network shown schematically in Figure 2C is implemented with *N* = 32 wedge neurons (based on anatomy^37^). For simplicity, we model the population of R5 neurons with a single variable *r_I_*(*t*) as in the sleep homeostasis model (section 2.1). The activities of wedge and R5 neurons are described by the following system of differential equations:

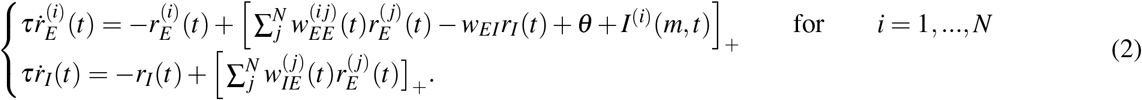

Here, *r_E_* (*t*) and *r_I_* (*t*) are the firing rates at time t of wedge and R5 neurons, respectively, *w_AB_* is the synaptic weight from population *B* to population *A*, *θ* is a constant background input onto wedge neurons, and *τ* is the effective population time constant. The matrix 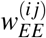 is initialized with a Gaussian function that depends on the distance between wedge neurons along the ring. The Gaussian has two parameters, the maximum amplitude, 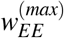, and the standard deviation σ (equation (38) in Methods). Figure 3A illustrates the connectivity from all the wedge neurons to wedge neuron 16. Additionally, *I^(i)^*(*m, t*) is an input from any modality to each wedge neuron *i* (for example (time-varying) visual or idiothetic input). This input encompasses input from ExR1 neurons, that process visual stimuli^18^, as well as from others populations. We assume that this input can be inhibited by dFB neurons, *r_S_*(*t*), as in the sleep homeostasis model (section 2.1), and is defined as a Gaussian function where the peak is located at a given wedge neuron *m* (equation (39) in Methods).

**Figure 3.**
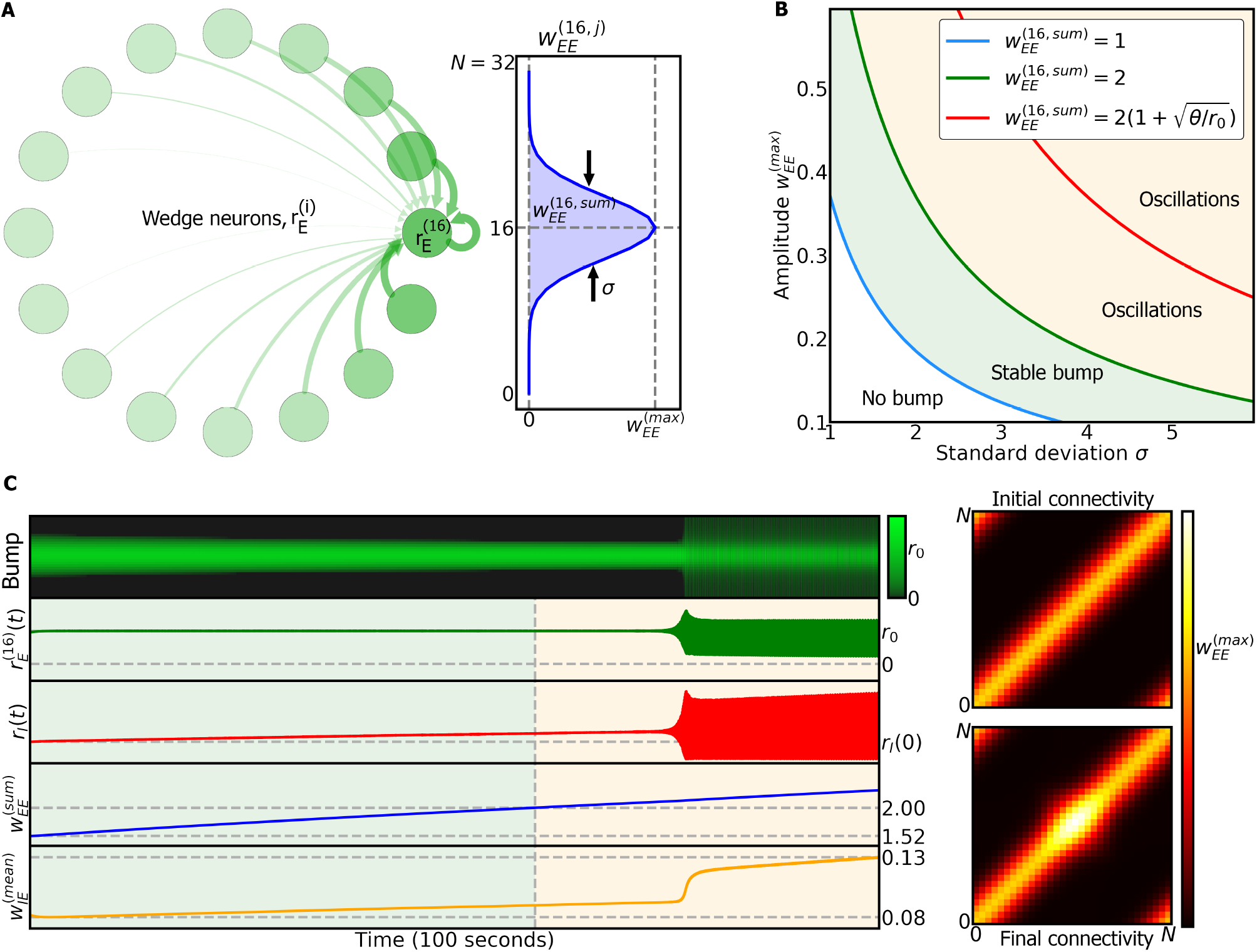
Dynamics of ring attractor network during wake phase. **A** Left side: representation of excitatory connections from all wedge neurons to wedge neuron 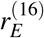. Right side: Gaussian connectivity from all wedge neurons to wedge neuron 16, with maximum amplitude 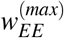 and standard deviation σ. **B** Model dynamics obtained in the fast-timescale limit (see Methods(4.7)) depend on the parameters of the excitatory connectivity, 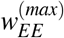 and σ. **C** Left side: dynamics during wake phase of the ring attractor model where the bump is located around wedge neuron 16. When the total excitatory weight, 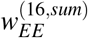, crosses a threshold, the bump starts to oscillate. Right side, top: initial excitatory connections between wedge neurons. Right side, bottom: final excitatory connections between wedge neurons; changes are a result of synaptic plasticity.

The inhibition of input to wedge neurons during sleep is motivated by the fact that self-motion inputs are not present, since the fly does not move during sleep. On the other hand, ExR1 neurons, which contribute to visual processing and locomotion, are inhibited by dFB neurons^18^. We hypothesize that other neural populations providing visual input to the ring attractor^23^ might require coincident activity from ExR1 neurons to reliably transmit visual information. This information might not be transmitted during sleep because of lack of activity from ExR1 neurons. This is consistent with the idea of an increased arousal threshold during sleep, where stronger stimuli are required to produce a behavioral response^38^.

The plasticity during the wake phase in recurrent connections between wedge neurons, 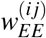, and from wedge to R5 neurons, 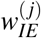 is modeled as follows:

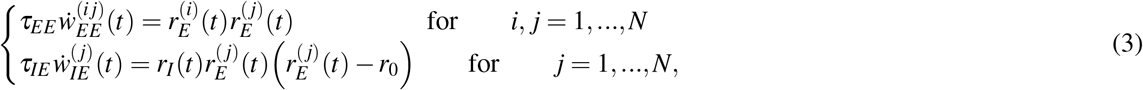

where *τ_EE_* and *τ_IE_* are time constants, and *r*_0_ is a positive presynaptic threshold. We assume that the dynamics of the plasticity rules are much slower than the dynamics of neural populations, so that *τ_EE_, t_IE_* ≫ τ, producing long-term plasticity. The synaptic weight between a presynaptic wedge neuron *j* and a postsynaptic wedge neuron *i* is represented by the time dependent matrix element 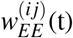. While the first plasticity rule is a linear Hebbian rule, the second is a triplet rule with presynaptic threshold; its behavior is similar to the linear Hebbian rule with presynaptic threshold, but it has a quadratic dependency on the presynaptic activity while ensuring no change in *w_IE_* if neural activity is zero (see derivation in Methods 4.5.1 and in^39^).

The plasticity rules are motivated by the assumption that the observed increase in activity and synaptic strength in R5 neurons^19^ balance the long-term potentiation in recurrent connections of wedge neurons, 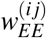. The plasticity rules therefore produce the following effects: (i) The recurrent synaptic connections between wedge neurons, 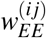, as well as the connections between wedge and ring neurons, *w_IE_*, get stronger during the wake phase. (ii) The firing rate of ring neurons *r_I_* increases during the wake phase. (iii) The amplitude of the bump in wedge neurons (which represents head direction) evolves always towards a constant setpoint, *r_E_* → *r*_0_. Note that activity is not constrained to the setpoint, but evolves towards it over time, since the setpoint is a stable fixed point for wedge neurons. Therefore, the bump amplitude will deviate from the setpoint due to any (for example visual or self-motion related) input (see Figure S4), consistent with experimentally observed behavior-related changes in bump amplitude^40^. These plasticity rules avoid the problem of a vanishing bump amplitude, as observed in the model with fixed connections (see Figure 2A and B).

How plasticity can drive the observed increase in R5 neuron activity with sleep pressure^19,27^ is currently unknown. Since it is the activity of R5 neurons, and not the plasticity, which is hypothesised to trigger sleep^18,19,27^, our models assume plasticity that directly modifies the activity of R5 neurons linearly (Figures 3C, 4A and 5A) (a possibility that is consistent with the data in^19^).

**Figure 4.**
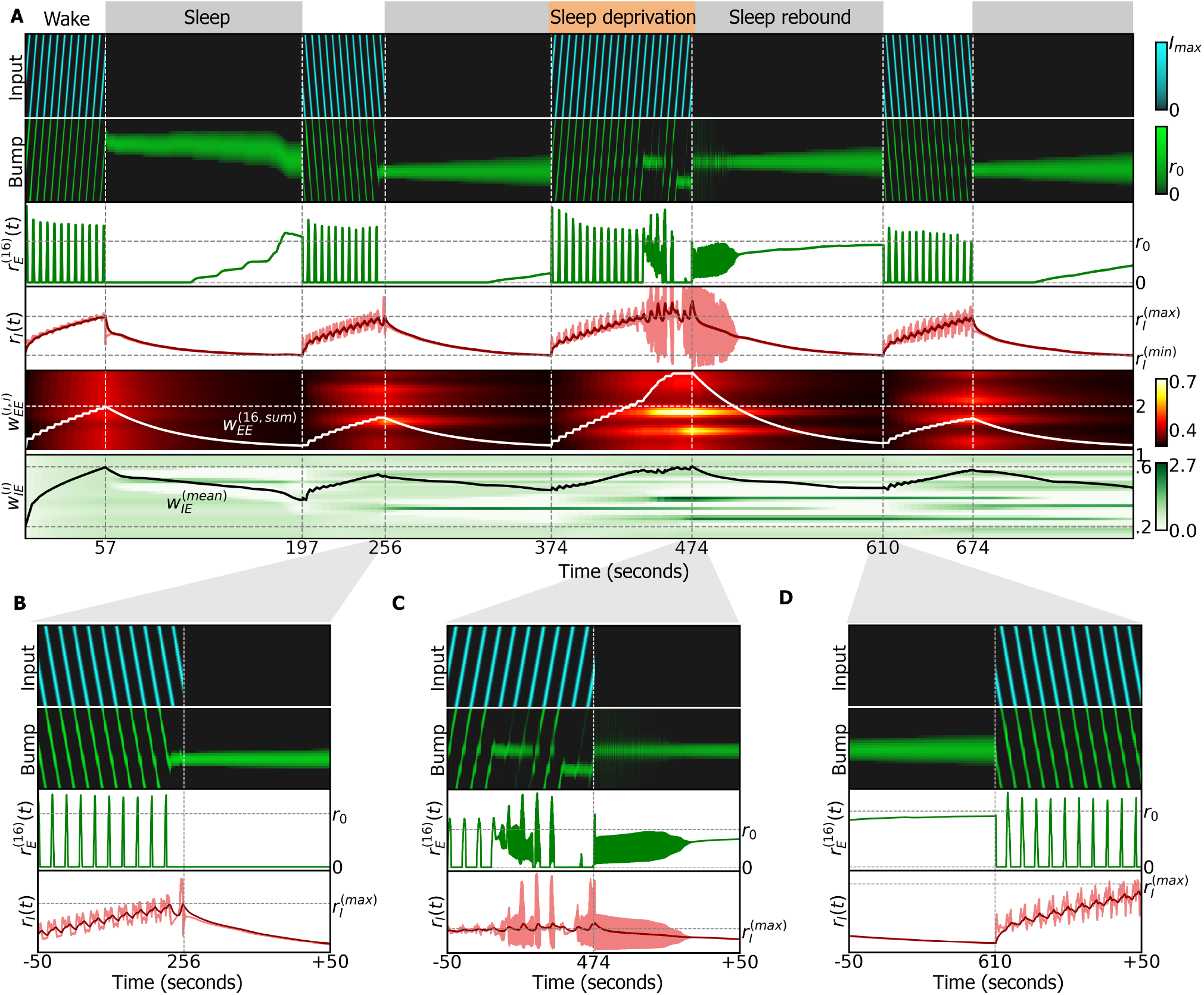
Simulation of ring attractor combined with sleep homeostasis model, using an exponentially decaying plasticity rule during sleep (equation (6)). **A** Entire simulation over a period of 800 seconds. White and grey regions indicate the sleep and wake phases, and correspond to dFB neurons switching off and on, respectively. Top row: input (inhibited during the sleep phase), alternating between clockwise and counter-clockwise rotations at 0.5H*z*. Second row: ring attractor bump activity. Third row: activity of wedge neuron 16. Fourth row, light red: activity of ring neurons. Dark red: filtered activity. Switching between sleep and wake is carried out by dFB neurons that switch on and off depending on filtered activity crossing thresholds 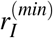 and 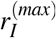. In the third wake epoch, sleep deprivation is produced by extending the inhibition of dFB neurons (*d*(*t*) = 1 during the orange top layout). Fifth row: diagonal elements of the connectivity matrix 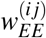. The white line is the sum of all excitatory connections to wedge neuron 16. It passes threshold 2 at around 240 seconds leading to oscillations. The full connectivity matrix 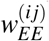 at the switch times is shown in Figure S7. Sixth row: connectivity 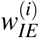; black line is the mean value. **B** Zoom-in around 256 seconds: switch from wake to sleep phase. **C** Zoom-in around 474 seconds: extended wake phase leads to oscillatory behavior. Circuit switches to sleep. **C** Zoom-in around 610 seconds: switch from sleep to wake phase.

**Figure 5.**
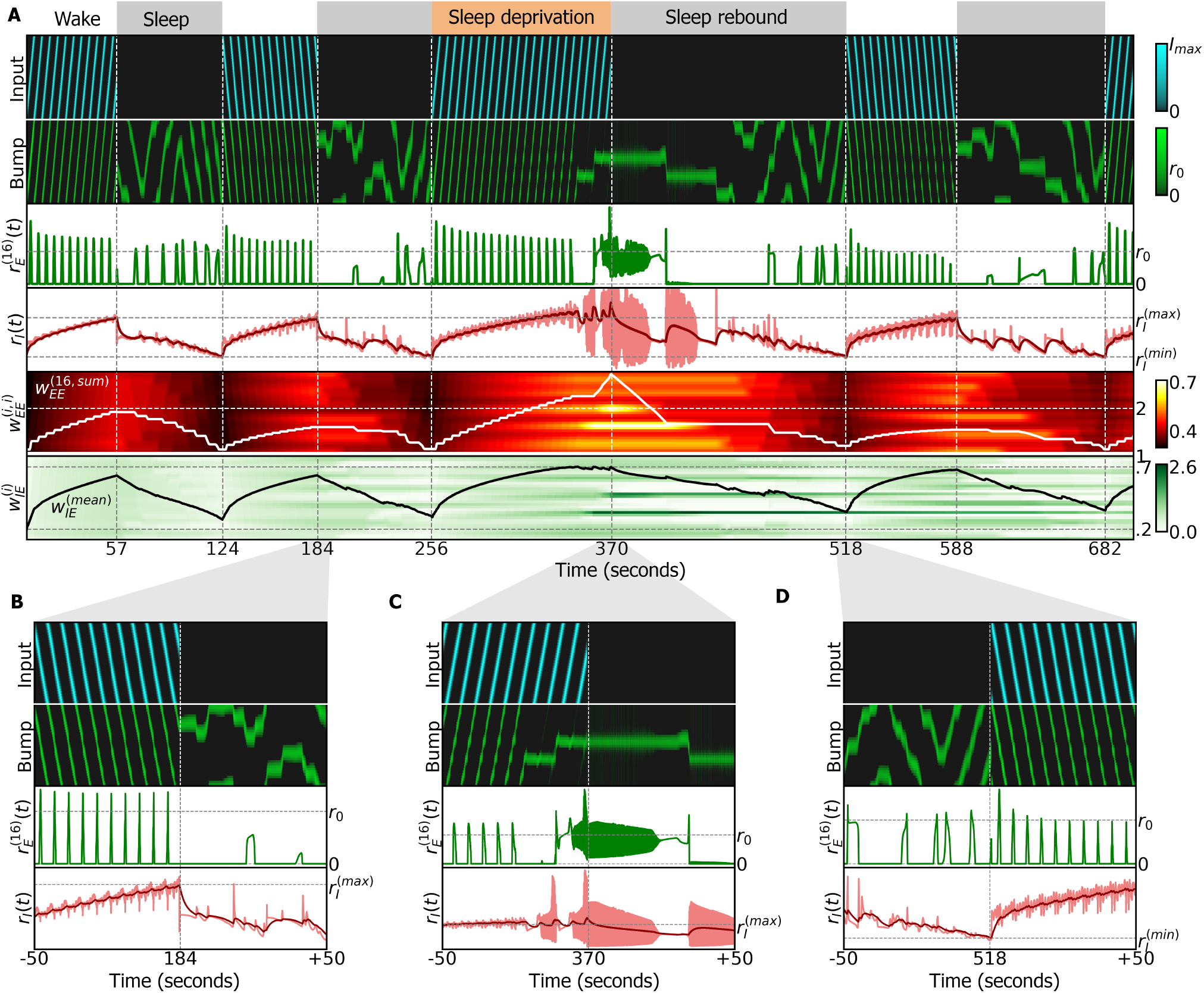
Simulation of ring attractor combined with sleep homeostasis model, using an anti-Hebbian plasticity rule during sleep (equation (7)). **A** Entire simulation over a period of 700 seconds, similar to Figure 4A. **B** Zoom-in around 184 seconds: switch from wake to sleep phase. **C** Zoom-in around 370 seconds: extended wake phase leads to oscillatory behavior. Circuit switches to sleep. **D** Zoom-in around 518 seconds: switch from sleep to wake phase.

A simplified model consisting of only one excitatory population representing wedge neurons and one inhibitory population representing R5 neurons is presented in Methods 4.5. Both populations are connected according to Figure 2C, including the plastic connections. This model shows overall similar characteristics to the full ring attractor network (see also next section).

### 2.5 Analysis of model stability during the wake phase

As shown below, the stability of a bump of activity in wedge neuron *i* is determined by its total excitatory input, 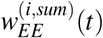 (Figure 3A). Here and in the following, we focus on wedge neuron 16, but the same analysis applies for all wedge neurons:

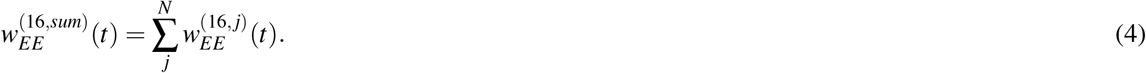

Figure 3B shows the different dynamic regimes of the bump as a function of the parameters 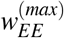 and s that determine the values of 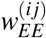 (see Methods (4.7) and Figure S6). The colored lines in Figure 3B are isolines of constant 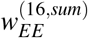, and correspond to the boundaries of distinct dynamics of the bump of activity in wedge neurons.

The boundaries are similar to the ones found in the simpler two-population model (Methods 4.5). The bump is stable around wedge neuron 16 if 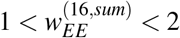. As the recurrent weights 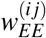 increase due to LTP during the wake phase, so do σ and 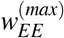. The bump starts to oscillate if recurrent connections are too strong, i.e. 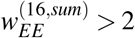. When 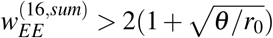, the bump keeps oscillating with very low activity in wedge neurons (see Methods 4.7).

Figure 3C illustrates the dynamics of the system with a bump centered in wedge neuron 16 (first row): as 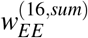 increases (fourth row) due to Hebbian plasticity, the weights 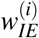 (represented by the mean) also increase (fifth row), leading to increased activity of R5 neurons (third row), which in turn maintains the amplitude of the bump in wedge neuron 16 constant (second row).

When 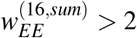, the bump starts to oscillate, as do R5 neurons (orange region in the last four rows). In addition, the plasticity rule in the recurrent connections 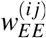 leads to an increase of the synaptic weights around the bump position (Figure 3C, right side, top *vs*. bottom).

### 2.6 Resetting the connectivity of the ring attractor during sleep

To limit synaptic growth as well as to avoid the instability of the bump and oscillations in R5 neurons, we introduce a sleep phase, as proposed by models of sleep homeostasis^18^. In this model, dFB neurons detect increased activity of R5 neurons and implement the switch between sleep and wake phases. In the model, we use a filtered version of R5 activity, which is motivated by the fact that R5 and dFB are not anatomically but functionally connected^19^. Additionally, this filtering prevents uncontrollable switching between phases in the oscillatory regime. The filtered activity of R5 neurons and the activity dFB neurons are modeled as follows:

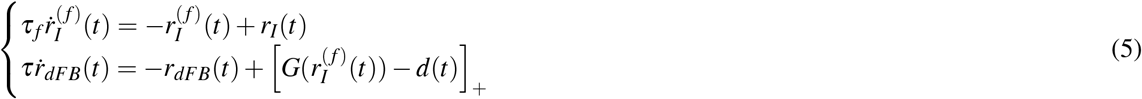

where τ*f* is the time constant of the low-pass filter (order of seconds); 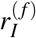, which we refer to as the switching signal, is a low-pass filter of activity in R5 neurons; and *r_dFB_*(*t*) is the activity of dFB neurons with its switching behavior modeled by equation (9). Finally *d* (*t*) is a variable which can produce sleep deprivation.

During the sleep phase, we assume that dFB neurons inhibit input to the ring attractor, similar to the sleep homeostasis model (see equation 39). In addition, our model resets the connectivity strength in 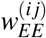 by a change in the plasticity rules during sleep. A simple way to restore the network connectivity 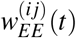 is to relax it to its initial values over the sleep phase:

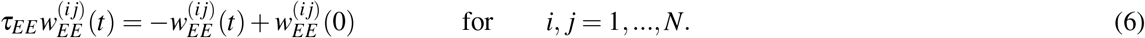

Note that we do not change the plasticity rule of 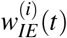, since this rule ensures 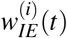 to always follow the trend (potentiation or depression) of 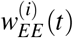 in order to maintain the bump amplitude at a setpoint. Figure 4A shows a simulation of the combined ring attractor and sleep homeostasis models ((section 2.1)) with plasticity. During wake phases (top, white region), dFB neurons are inactive (*r_dFB_*(*t*) = 0, not shown), and during sleep phases (grey region) dFB neurons are active (*r_dFB_*(*t*) = 1). In the wake phase, a rotating input with a constant frequency of 0.5*Hz* is provided; the input reverses direction between consecutive wake phases (top row). During the wake phase, the bump in the ring attractor closely follows the input (second row), while the activity of R5 neurons increases (light red line in the fourth row). The second to last row shows the diagonal elements of 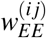 while the last row shows 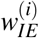. When the switching signal reaches the upper threshold, 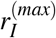, dFB neurons switch the model to the sleep phase, where the plasticity rule in 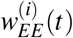 changes and input to wedge neurons is inhibited (Figure 4B). During sleep, the bump in wedge neurons stays in place, maintaining the last head direction of the fly before the sleep phase while the activity of R5 neurons slowly decreases. As with the two-population model (Methods 4.5), the timescale of sleep and wake phases depends on the time constants of the plasticity rules.

Once the switching signal reaches the lower threshold, the system switches back to wake phase and the input is turned on again (Figure 4D). Furthermore, if we prevent dFB from switching to the sleep phase (by setting the variable *d*(*t*) = 1) and thus extend the wake period (sleep deprivation, upper orange region), 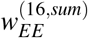 for wedge neuron 16 crosses the boundary for stability, 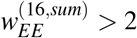. In this case, the bump in the ring attractor starts to oscillate (Figure 4C; see section 2.5). Towards the end of the extended wake phase, the bump stops tracking the input. In the subsequent sleep phase, more time is required to reset the excitatory weights and to reach the lower threshold 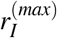, resulting in sleep rebound (Figure 4A).

### 2.7 Resetting the ring attractor during sleep using autonomous dynamics

An alternative mechanism for the network to be reset during sleep is an anti-hebbian plasticity rule in 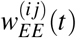, such as the following:

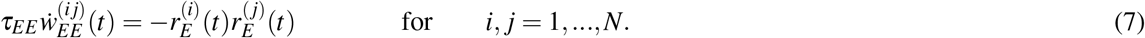

This rule produces LTD with correlated activity between neighboring wedge neurons. Figure 5A shows a simulation of this model. In the wake phase, the behavior is, as expected, the same as in the model without anti-hebbian plasticity (Figure 4A). During sleep, the autonomous bump movement resets synaptic connections and the activity of R5 neurons decreases. Heterogeneity in the weights 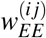 (due to synaptic plasticity) makes the bump drift across wedge neurons. These autonomous dynamics reset the connectivity in the network^7^. The amplitude of the bump during autonomous dynamics is at setpoint level, that is, below the activity level resulting with input to the attractor network.

Wake phases without continuous input can also show drift (see Figure S8 with intermittent input). This wake drift is however different from sleep drift, since it ends once the bump reaches the location of strongest recurrent excitation, making the synaptic weights grow in this location until sleep is initiated. Such wake drift can be reduced in our model by slowing the plasticity rules and with close to homogeneous coverage of the bump movement across all wedge neurons (see Figure S9 and Methods (4.9)).

To investigate how such autonomous dynamics of the bump during sleep are linked to the dynamics during the preceding wake phase, we provided sinusoidal inputs a range of amplitudes *A* and frequencies *f*. In Figure 6A, we show a simulation with fixed amplitude, *A* = *N*/4 and different frequencies in each wake phase (0.1,0.5, and 1 Hz). During sleep, the bump revisits wedge neurons that were active in the preceding wake phase, as seen in the distributions of the time spent around each wedge neuron during the first (left), second (center) and third (right) wake phase (in blue) and during the following sleep phase (in grey) (Figure 6B). To further probe the amplitude dependence, we simulated a wake phase and the subsequent sleep phase, and in each simulation varied the amplitude *A* of a sinusoidal input in the range [0, 16], with fixed frequency of *f* = 1*Hz*, during the wake phase. The standard deviation (STD) of the paths of the bump during the wake (blue) and sleep (grey) phase closely match (Figure 6C). Similarly, we investigated the frequency dependence with a stimulus with fixed amplitude, *A* = *N*/4, and varying frequency in the range [0.1,1.5]*Hz*. The number of oscillatory cycles grows linearly with the input frequency in the wake phase (blue), but remains constant during sleep. Therefore, the dynamics during sleep do not depend on the input frequency during the wake phase (Figure 6C). Additionally, the frequency of oscillations of the bump around the ring can increase as the bump approaches the lower threshold of the switching signal (before waking up, Figure S9A). Such autonomous dynamics are reminiscent of activity observed during sleep in mice^41^.

**Figure 6.**
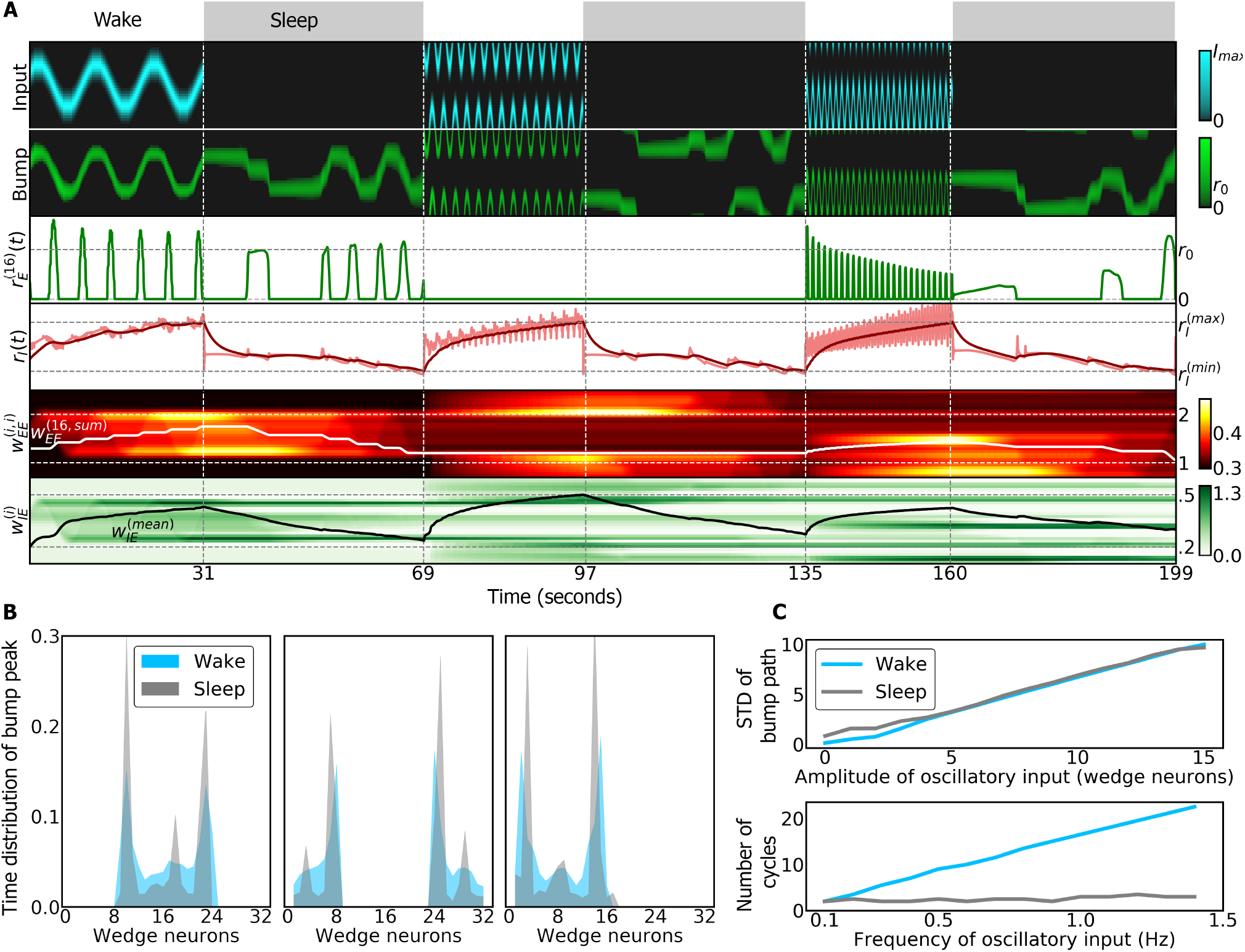
Simulation of the ring attractor network with sinusoidal inputs during wake phase. **A** In each wake phase, a sinusoidal input is provided to the ring attractor (top row) with increasing frequencies in consecutive wake phases. During sleep, an autonomous bump of activity revisits wedge neurons active in the previous wake phase (second row). The third row shows the activity of wedge neuron 16 and the fourth row shows the activity of the ring neuron (light red) and its filtered activity (dark red) used to switch between wake and sleep phases. The last two rows represent the synaptic weights that increase during wakefulness and decrease during sleep. **B** Normalized distribution of time that the bump peak is localized in each wedge neuron during wake (blue) and sleep (grey) phases. The first, second and third plots show the distribution for the first, second and third wake and sleep phases, respectively. **C** Top: standard deviation (STD) of the bump path during a total of 15 simulations where different amplitudes of a sinusoidal inputs are provided during the wake phase. During the wake phase, the STD of the bump path is proportional to the amplitude (grey). During sleep, the autonomous bump path has a correlated STD (Pearson correlation coefficient: 0.99 (*p* = 1.5 · 10^-17^). Bottom: number of cycles of the bump path during a total of 15 simulations with different frequencies of a sinusoidal input during wake phase. During the wake phase, the number of cycles is proportional to the input frequency. However, during sleep, the number of cycles does not correlate with the input frequency: Pearson correlation coefficient: 0.76 (*p* = 0.0015).

## 3 Discussion

In the brain of *Drosophila*, structurally similar neurons in the center of the brain have been assigned functionally very different roles. On the one hand, navigation related ring neurons encode spatial memory or visual features^22,23,42^. For these ring neurons, ring attractor networks offer a compelling structure-function relationship which can provide a rationale for their ring-shaped morphology. On the other hand, sleep related ring neurons serve as homeostatic sleep integrator, encoding sleep drive through structural^19,28^ and activity changes^19,27^. The connectome additionally shows multiple interactions between these sleep and navigation related circuits^14,30^.

To elucidate the relationship between these navigation and sleep functionalities of ring neurons, and to address how the head direction system can operate in the face of plasticity in connected circuits, we therefore asked what role the homeostatic integrator could play in the ring attractor framework.

To address this question, we used the sleep homeostasis model proposed in^18^ as a starting point. The connectome shows that this circuit is not isolated but interacts with the head direction system (Figure 2A). When connectivity in this circuit is fixed, however, the increasing activity of R5 neurons (which encode sleep drive) decreases the amplitude of the bump of activity in the ring attractor (Figure 2D). To overcome this problem of vanishing activity, we therefore propose a model with plasticity in R5 neurons (which is experimentally observed) and with hypothesized recurrent plasticity between wedge neurons. In this model (Figure 2C), sleep drive balances plasticity in wedge neurons, which are now able to maintain a bump of activity that evolves towards a constant amplitude setpoint over time. The model also allows variability in the bump amplitude with external (for example visual or self-motion related) input (Figure S4), consistent with experimentally observed behavior-related changes in bump amplitude^40^.

However, prolonged activity during wakefulness ultimately leads to unstable behavior (oscillations in Fig. 3C and 5A). Therefore, to restore the connectivity in the head direction system to baseline, we introduced a sleep phase, in agreement with models of sleep homeostasis^18^, where the synaptic connections between wedge neurons are reset by LTD. While the time course and dynamics of this reset is not known, we here investigated two alternatives. In one case, while dFB neurons inhibit input, the ring attractor resets to its initial state while the bump stays in place. In the second case, an anti-Hebbian rule resets the ring attractor with autonomous dynamics. These dynamics are linked to the dynamics during wakefulness through their spatial (Figures 6A and 6B) but not through their frequency distributions (Figures 6A and 6C). The amplitude of the bump during autonomous dynamics is at setpoint level, that is, at the level of activity to which the amplitude settles in the absence of (visual or idiothetic) input.

In the proposed model, heterogeneities in recurrent connections of wedge neurons can also lead to drift during the wake phase with intermittent inputs (see Figure S8). While this is consistent with the heterogenities observed in the connectome (Figure S1B), we minimize drift in the model by the slow dynamics of the plasticity rules together with assumed homogeneous activation of wedge neurons over time S9). Other solutions to avoid drift in ring attractors have however been developed (for example^43-45^).

Many aspects of this model can also be captured by a simpler two-population model, which shows similar dynamics and related boundaries between the different dynamic regimes (Figures S3B and 3B).

The introduction of plasticity was motivated by the observation of structural, synaptic and functional changes in R5 neurons^19,27,28^ as well as their interaction with the head direction system as suggested by the connectome^14,30,46^. The proposed combined sleep homeostasis and ring attractor model can capture this increase in activity in R5 neurons during wakefulness^19,28^ (Figures 6 and 5). Additionally, in the proposed models, sleep deprivation leads to a qualitative change in the behavior of R5 neurons towards oscillatory dynamics which is reminiscent of the experimentally observed transition to bursting dynamics^19^ or increase in oscillatory dynamics^27^. Whether sleep deprivation compromises the head direction system in behaving flies is currently not known, although navigation related memories are for example affected by sleep deprivation in bees^47,48^.

The proposed model makes several predictions that could be tested experimentally. First, we assume LTP in the recurrent connections of wedge neurons during wakefulness (which could also be achieved through an intermediate population^34^). This plasticity in *w_EE_* could be a result of correlated activity between neighboring wedge neurons. Second, as a result of this plasticity, the bump width changes with *w_EE_*, decreasing over time spent awake due to LTP in wedge neurons and therefore increasing spatial resolution (and *vice versa* during sleep, Figure S6C). Generally, there is a range of bump widths that can be sustained by the ring attractor (Figure 3). A third model prediction is that extended wakefulness can disrupt the head direction system by producing oscillatory or bursting behavior and will lock the bump position in place (independent of external input, Figure 5A, sleep deprivation). Additionally, sleep results in autonomous dynamics in the ring attractor model (Figure 5), with the network transitioning towards faster dynamics towards the end of the sleep phase (Figure S9). Such autonomous dynamics are reminiscent of activity of the head direction system observed in mice during sleep^41^.

We further assume that the plasticity rule in the head direction system changes during sleep from LTP to LTD. This change of plasticity has been proposed in several models of sleep (for example^49^; see^4^ for review). A potential mechanism could for example be neuromodulation of an STDP gate^50,51^, which has been observed in insects^52,53^ and could involve the strong innervation of the central complex by neuromodulatory neurons^54^. For example, ExR1 neurons described in^18^ (modeled in Figure 1A and B) could produce the switch in plasticity between sleep and wake phases, potentially through neuromodulation (similar to the related serotonergic ExR3 neurons^30,55^).

The resulting weakening of synaptic strength during sleep underlies several hypotheses about sleep function^4-6,8^. The approach implemented here is based on the idea of reverse learning^5-7^: during sleep, attractors within the ring attractor network generated during wakefulness are removed and the corresponding increased weights are weakened. Autonomous dynamics during sleep could be functionally relevant for memory consolidation and organization^56^. For instance, flies could partially replay (in wedge neurons) trajectories during sleep that they performed during navigation in the wake phase (see Figure 6A) which could be used by downstream circuits to consolidate navigation related memories. Navigation memories are for example consolidated during sleep in bees^47^, and replay of neural activity in the central complex during sleep has been suggested to consolidate courtship memory in flies^57^.

While synaptic changes during sleep and wakefulness are observed across the fly brain (for example^58^), one could hypothesize that such activity related changes are stronger in areas where activity is persistent with a possible role in working memory, such as the head direction system^25,42^. Therefore, inhibitory R5 neurons might increase their activity faster and require resetting through sleep sooner than other transiently active neurons, ultimately being responsible for signaling sleep drive. We additionally did not differentiate between different ring attractor inputs (for example visual or idiothetic) and such different signals could also be integrated in different ring neurons or homeostats^11^ (taking for example into account that visual experience increases sleep need^59^).

The connectome shows that both the head direction as well as the sleep homeostasis circuits encompass a large number of connected cell types in the central complex^30,60^. Nevertheless, strongly simplified models of ring attractor networks with only a limited subset of actually involved cell types have proven useful for the description of the head direction system. Similarly, for the sleep homeostasis circuit, many more connected cell types could be considered and we here only investigated a simplified network that nevertheless can capture several experimental observations.

Overall, the interaction of the homeostatic integrator and the head direction systems together with mounting evidence for a close structure-function relationship in these circuits, suggest that a relationship between the control and function of sleep could be established in this network using theoretical modeling and experiments.

## Acknowledgements

We would like to thank Raoul-Martin Memmesheimer for helpful discussions and comments on the manuscript, and Gerry Rubin, Julijana Gjorgjieva, Felipe Kalle Kossio, and Marina Elaine Wosniack for helpful discussions.

## 4 Methods

### 4.1 Anatomy based on the fly connectome

The connectivity of the proposed model is based on the fly connectome^14^ and incorporates the populations R5 (ER5), ExR1, EPG and EL, as described in the Neuprint database. Each population and its innervation in the ellipsoid body are shown in Figure S1A. EL and EPG neurons, respectively, have diagonal connectivity matrices within and across populations as shown in Figure S1B.

### 4.2 Numerical simulation of models

We numerically solved all models with forward Euler with a time step of *dt* = 0.0001 seconds. Our code is implemented in Python, and we will make it available upon publication.

### 4.3 Sleep homeostasis circuit

In this and the following, we use rate-based models to simulate dynamics of entire neural populations and dynamics of single wedge neurons. The differential equations used to model the sleep homeostasis circuit are as follows:

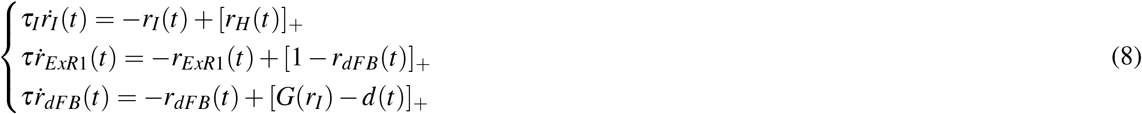

The switch behavior of dFB neurons is modeled by simple hysteresis, according to the following equation:

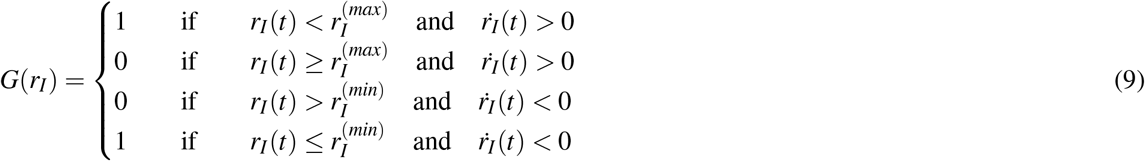

Such a switch behavior in dFB neurons could be implemented, for example, by adding an additional wake-promoting population, which together with dFB neurons, could mutually inhibit each other to create a flip-flop switch, similar to sleep models proposed in mammals^61-64^. Candidates for the wake-promoting population in the fly are dopaminergic neurons in the PPM3 and PPL1 clusters^29^. Alternatively, this switch behavior could be generated by a single-cell mechanism in dFB neurons, which are known to increase excitability with extended wake time^29^.

In this model, the wake and sleep time depend on the effective time constant *τ_I_* and the thresholds 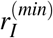 and 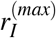. The time spent in the sleep phase, *t_S_* (sleep time) as a function of the time spent in the wake phase, *t_W_* (wake time), can be computed by solving the differential equation for R5 neurons, *r_I_*, during the sleep phase and wake phase, respectively:

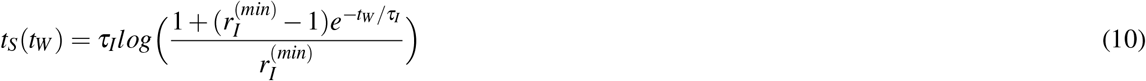

Considering that *t_W_* is small, we can expand this expression in a Taylor series, taking only first order terms:

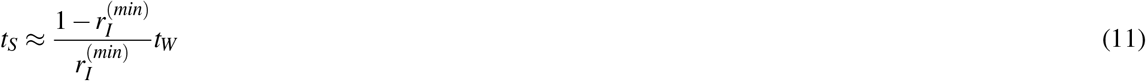

For small waking time periods, the sleep time *t_S_* increases linearly, resulting in sleep rebound as required for homeostasis (increased time spent awake leads to more sleep afterwards). However, for long wake times, the preceding sleep time saturates at a constant value 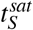, given by the following expression:

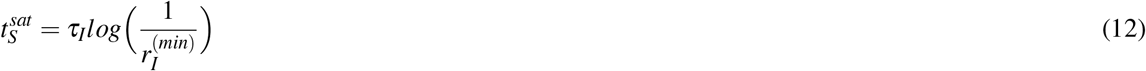

This saturation time prevents very large sleep times after large preceding wake times, and it is also a feature observed experimentally^65^.

### 4.4 Sleep homeostasis and ring attractor with fixed connections

We asked how increasing activity in R5 neurons affects the head direction circuit in the absence of plasticity. Given the fact that R5 and wedge neurons are connected, we modeled a ring attractor network where wedge neurons encode head direction and R5 neurons provide increasing inhibitory input to wedge neurons. The model is shown schematically in Figure 2A and described by the following system of equations:

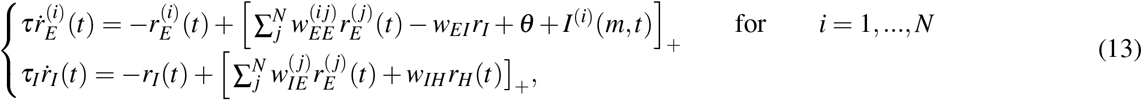

where *r_E_*(*t*)^(*i*)^ represents the activity of a wedge neuron *i* (in total, *N* = 32) and and *r_I_*(*t*) is the population activity of R5 neurons and *τ_I_* accounts for slow dynamics as in the phenomenological sleep homeostasis model (section 2.1). We only model the wake phase for simplicity, where dFB neurons are assumed to have zero activity and activity in ExR1 neurons is defined as *r_H_* (*t*) = 1, similar to the sleep homeostasis model. We neglect the connection from wedge to ExR1 neurons for simplicity, since we focus on the interaction between wedge and R5 neurons. The weights *w_AB_* represent the connectivity from population *B* to population *A*. The recurrent connectivity 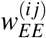 is a matrix, in which for a given presynaptic wedge neuron, *i*, the element (*i j*) is given by a gaussian function that depends on the distance to the postsynaptic wedge neuron *j* along the ring, given by equation (38) (see for example Figure 3C, left top). In this model, the values of the synaptic weights *w_IE_* and *w_IH_* were chosen such that the activity of ring neurons increases^19^.

Figure 2B shows a simulation of the model, where a rotating input *I*(*m,t*) is provided to wedge neurons at 0.5 Hz (top row, blue). The activity of R5 neurons increases, as imposed by our parameter choice (third row, red). The wedge neurons, (second row, green) follow the rotating input while receiving this increasing inhibition, such that the bump amplitude decreases over time until inhibition gets strong enough so that the bump vanishes.

### 4.5 Two-population model with plasticity for R5 and wedge neurons

To simplify the analysis and build intuitions about the complete ring attractor model combined with the sleep homeostasis circuit, we first developed a simpler model. This model is a population model based on an excitatory-inhibitory network^66^ (Figure S3A) and describes the interaction between wedge and R5 neurons. The dynamics of (excitatory) wedge and (inhibitory) ring neurons are described by the following system of differential equations:

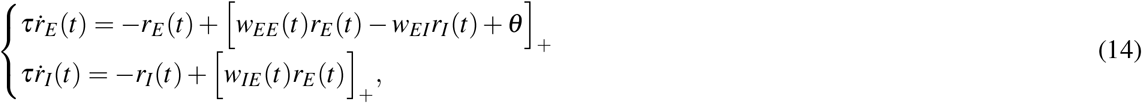

where *r_E_* (*t*) and *r_I_* (*t*) are the firing rates at time *t* of wedge neurons and ring neurons, respectively, *w_AB_* is the synaptic weight from population *B* to population *A*, *θ* is a constant background input onto wedge neurons, [·]_+_ is a threshold-linear function to ensure positive-valued firing rates, and *τ* is the effective population time constant.

#### 4.5.1 Plasticity rules

We additionally introduce plasticity rules for the excitatory weights *w_EE_* and *w_IE_* during the wake phase (which we motivate below):

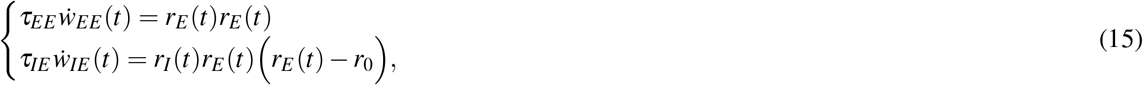

During the sleep phase we change the plasticity rule in *w_EE_*, while leaving unchanged the plasticity rule in *w_IE_*:

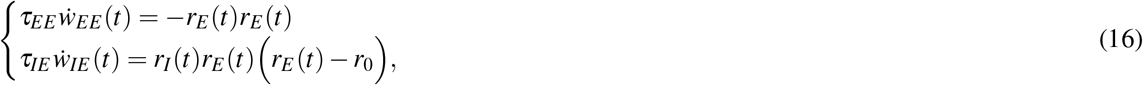

where *τ_EE_* and *τ_IE_* are time constants, and *r*_0_ is a positive presynaptic threshold. While the first plasticity rule is a linear Hebbian rule, the second is a triplet rule with presynaptic threshold. These plasticity rules can be extracted from a general form of Hebbian plasticity. A general hebbian plasticity rule for a synaptic weight *W_ij_* can be defined as follows:

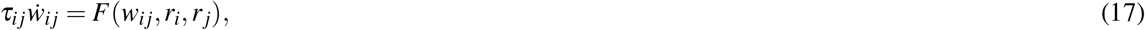

where *τ_ij_* is the time constant of the rule, and *F*(·) is a function that depends on the synaptic weight, *W_ij_*, and on pre- and postsynaptic activities, *r_j_* and *r_i_*, respectively^39^. The function *F*(·) needs to fulfill Hebb’s condition: to produce a change in the synaptic weight *W_ij_*, the pre- and postsynaptic neurons must be active: *r_i_* > 0, *r_j_* > 0. In principle, this function is unknown, but we can expand it in a Taylor series^39^ around *r_I_* = *r_E_* = 0:

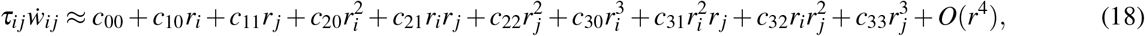

where each coefficient depends on the connection strength *c_mn_ = c_mn_*(*w_ij_*). The values of these coefficients determine the plasticity rule. For instance, Hebbian plasticity rules that are linear in the neural activities can be obtained by setting second or higher order coefficients to zero^39^. Keeping higher order coefficients leads to rules with non-linearities.

We assume that the plasticity rule in *w_EE_*(*t*) during the wake phase is linear, obtained by setting *c*_21_ = 1 and all other coefficients to zero, while the plasticity rule in *w_IE_* is non linear on the presynaptic neural activity, obtained by setting *c*_21_ = −*r*_0_, *c*_32_ = 1 and the other coefficients to zero (equation (16)). To ensure that synapses remain excitatory or inhibitory throughout the system’s dynamics at any time, the plasticity rules are threshold-rectified at zero if the synaptic weights are zero:

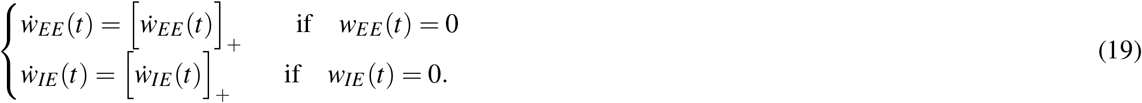

These plasticity rules are motivated by the fact that R5 neurons produce sleep drive by increasing activity and synaptic strength^19^ for balancing LTP in wedge neurons. The plasticity rules in equation (16) produce the following effects: *(i)* The synaptic connection between wedge neurons, *w_EE_* and between wedge and ring neurons, *w_IE_*, get stronger during the wake phase. *(ii)* The firing rate of ring neurons *r_I_* increases during the wake phase. *(iii)* The firing rate of the entire population of wedge neurons evolves always to a constant setpoint, *r_E_* → *r*_0_.

**Table S1.** Parameter values used in the two-population model. Time constants *τ*, *τ_f_*, *τ_EE_* and *τ_IE_* are measured in seconds. We consider the rest dimensionless quantities.

| $\tau$ | $\tau_f$ | $\tau_{EE}$ | $\tau_{IE}$ | $w_{EI}$ | $\theta$ | $r_0$ | $r_I^{(min)}$ | $r_I^{(max)}$ |
| --- | --- | --- | --- | --- | --- | --- | --- | --- |
| 0.01 | 2 | 10000 | 10000 | 0.5 | 10 | 10 | 20 | 35 |

Finally, we note that the dynamics of the plasticity rules are much slower than the dynamics of neural populations, so that *τ_EE_, τ_IE_* ≫ *τ*. The parameters for the two-population model used for simulations are shown in Table S1, but the following stability analysis is performed without any assumption on the parameter values.

#### 4.5.2 Stability of the two-population model

##### Fast-timescale limit

In the fast-timescale limit, we can assume that *τ_EE_*, *τ_IE_* → ∞, meaning that synaptic plasticity is sufficiently slow compared to the dynamics of the neural populations so that it can be assumed to be constant. This offers the advantage of treating the synaptic weight *w_EE_* as a free parameter with a fixed value, assuming that *w_IE_* has already evolved through its plasticity rule to its equilibrium value, *r_E_* → *r*_0_. Therefore, for a given value of *w_EE_*, we set the value of *w_EI_* such that the fixed point for the wedge neurons is *r*_0_. In this way, the value of *w_IE_* is coupled to the value of *w_EE_*. The stability of the 2-dimensional system given by equations in (14) is then analyzed with respect to the value of *w_EE_*. Since the system is piecewise linear due to the threshold function [·]_+_, we perform a linear analysis assuming that the inputs to the neurons are positive. Under these conditions, the fixed point of the system, 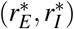, is given by the following expressions:

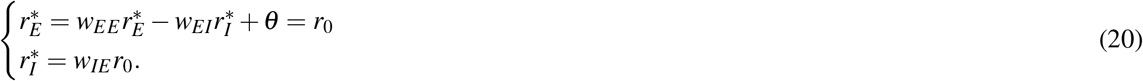

Since we force the fixed point of wedge neurons to be *r*_0_, we can extract the equilibrium value of *w_IE_* as a function of *w_EE_*:

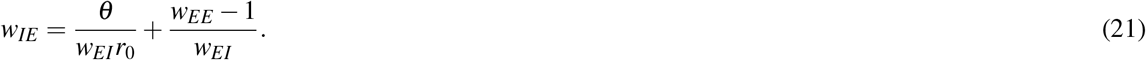

The fixed point of the system can be described with respect to *w_EE_* as:

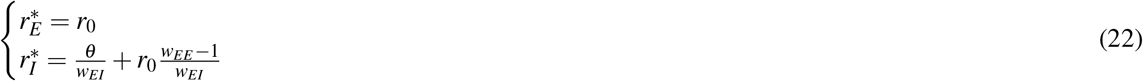

Both the fixed point of ring neuron activity, 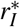, and the equilibrium value of the connectivity, *w_IE_*, depend linearly on *w_EE_*, implying that if *w_EE_* increases, both *w_IE_* and 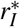 increase as well as long as the fixed point is stable. We analyze the stability of the system by calculating the eigenvalues:

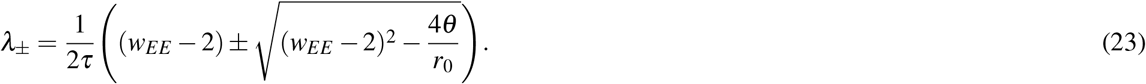

Both eigenvalues are shown in Figure S2A with respect to different values of *w_EE_*. This leads to four different cases:

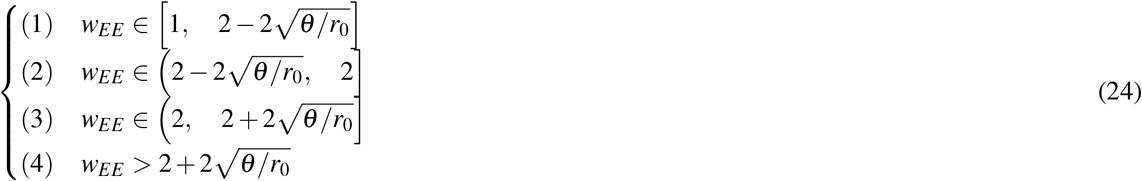

In case (1), both eigenvalues are real and negative: the fixed point is stable (equation (22)). In case (2), the real part of the eigenvalues is negative and the imaginary part is non-zero: the system evolves towards the fixed point with damped oscillations. In case (3), the real part of the eigenvalues is positive and the imaginary part is non-zero: the system diverges towards infinity, oscillating with amplitudes which increase exponentially. In case (4), the eigenvalues are real and positive: the fixed point is unstable. This analysis predicts a bifurcation in the stability of the fixed point when *w_EE_* = 2. This behavior is shown in Figure S3C.

The non-linearity of the linear threshold function changes the behavior of the model slightly. While the behavior stays the same for the cases (1), (2) and (4), because the model is mostly in the linear regime, case (3) differs and the non-linearity produces stable cycles around the fixed point. This behavior is found empirically from simulating the non-linear model, and is summarized in Figure S3B.

##### Slow-timescale limit

In the slow-timescale limit we consider that the firing rates change sufficiently fast compared with the synaptic weights so that these changes can be considered instantaneous (*τ* → 0). We therefore analyze the conditions under which the synaptic rules in equations (15) and (16) stabilize the model. We again first consider the linear range of the function [·]_+_ where the inputs to the neurons are positive. We approximate the instantaneous value of the firing rates in equation (14) as follows:

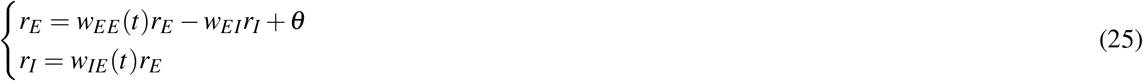

This linear system allows extracting the values of *r_E_* and *r_I_* in terms of the synaptic weights as:

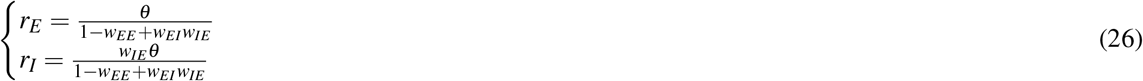

We can now compute the vector field for wedge and ring neuron activity as a consequence of the slow dynamics of synaptic plasticity:

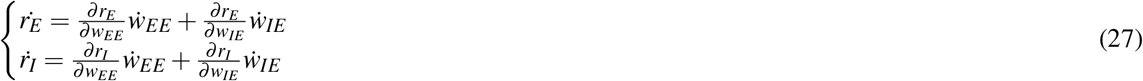

#### 4.5.3 Two-population dynamics during wakefulness

Considering the plasticity rules during the wake phase (15), equation (27) leads to the following system of differential equations:

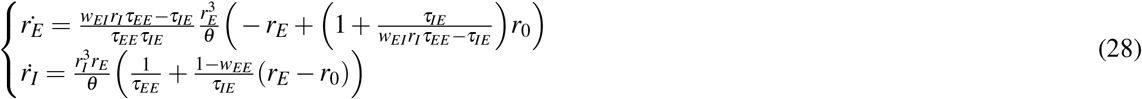

The second equation gives the dynamics of ring neurons, which increase activity with 1 /*τ_EE_*. The first equation gives the dynamics of the population of wedge neurons approaching a setpoint only when the effective decay time constant (the first factor in the right hand side) is positive, otherwise the equation diverges to infinity and the system is unstable. This gives the following criterion for *τ_EE_* and *τ_IE_*:

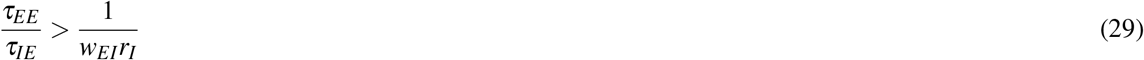

This relationship supports the idea that plasticity in *w_IE_* has to be fast enough with respect to the LTP in *w_EE_*; otherwise, if *w_EE_* increases faster than *w_IE_*, the model diverges. Let us compute the upper limit of inequality (29), which corresponds to the minimum of *r_I_*. For that, we approximate the firing rate of wedge neurons by *r_E_* = *r*_0_, and *r_I_* = *w_IE_r_0_*. Then, we can write *w_IE_* as a function of *w_EE_* as in equation (21). The upper limit of inequality (29) will therefore happen at the minimum of *w_EE_*. As the minimum is *w_EE_* = 1, the upper limit of the stability condition is:

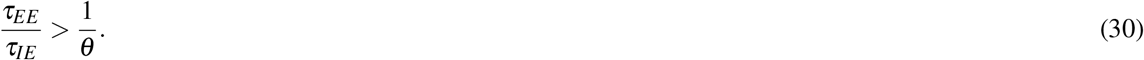

When this condition holds, the setpoint of wedge neurons in equation (28), which is not *r*_0_ as approximated previously, is given by

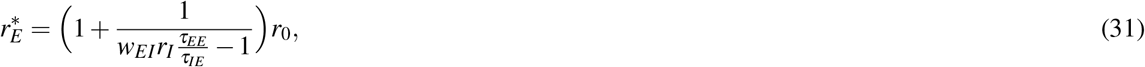

which is generally different from *r*_0_ due to the inertia of the dynamics of *w_EE_*. In the limit *τ_EE_* → ∞ (no plasticity in *w_EE_*), the fixed point is *r*_0_, as expected from the fast-timescale analysis.

Figure S2B shows the vector field for the system of equations (28). The green line shows the trajectory of the setpoint in wedge neurons as the activity of ring neurons increase. As *r_I_* increases due to increasing *w_EE_*, the setpoint in *r_E_* approaches *r*_0_.

During wakefulness, the fixed point for wedge neurons 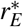 remains mostly constant, and the fixed point for ring neurons 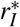 changes with *w_EE_* (Figure S3B). If *w_EE_* ≤ 2 (light green region), the fixed point 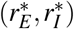 is stable, and both neural populations evolve towards these values. With increasing *w_EE_*, the fixed point of ring neurons 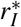 also increases while the fixed point of wedge neurons 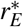 remains constant. If *w_EE_* increases further to 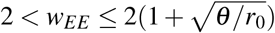, the model enters a regime of stable oscillations (light orange region of Figure S3B). In this regime, both neural populations oscillate around the fixed point with a frequency that changes with *w_EE_* (see Figure S2A, as explained in the fast-timescale limit. In addition, ring neurons increase their amplitude of oscillations as *w_EE_* increases. Finally, when 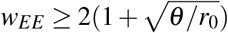 (light red region), the fixed point is unstable and the activity of both populations diverges.

Figure S3C illustrates the dynamics of the full system, i.e. the activity of wedge and ring neurons, *r_E_* (*t*) and *r_I_* (*t*), as well as synaptic weights, *w_EE_*(*t*) and *w_IE_* (*t*). In the beginning (light green region), the fixed point of ring and wedge neurons is stable because *w_EE_* < 2. When this boundary is crossed, the system enters the regime of stable oscillations (light orange region). Also, in the stable region of the simulation in Figure S3C, *w_EE_, w_IE_* and ring neuron activity *r_I_* increase, while the activity of wedge neurons *r_E_* remains constant as imposed by conditions (i)-(iii). *w_EE_, w_IE_* and *r_I_* constitute therefore a measure of how far the network has moved from its initial state.

#### 4.5.4 Two-population model dynamics during sleep

In order to reset the system back to its stable state (*w_EE_* < 2) after prolonged activity (wakefulness), we introduce a sleep phase with inverted plasticity^5-7^. For this we assume that during sleep the recurrent connection between wedge neurons, *w_EE_*, gets weaker through LTD^4,7^ while the plasticity rule for *w_IE_* is the same as in the wake phase (equation (16)).

We can perform the same analysis during sleep by considering the plasticity rule in equation (16) during sleep so that the equations (27) lead to the following system of differential equations:

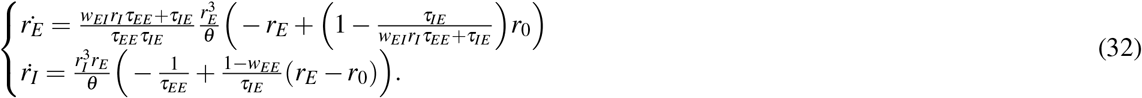

The second equation shows how ring neurons decrease their activity with 1 / *τ_EE_*, at the same rate as in the wake phase. The first equation shows a fixed point for wedge neurons that is lower than *r*_0_, due to the inertia of a decreasing *w_EE_* during sleep, given by:

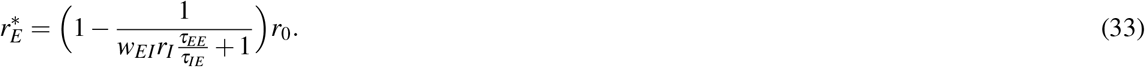

Figure S2C shows the vector field given by equation (32) during the sleep phase, where the trajectory of the setpoint of wedge neurons (the green line) diverges from *r*_0_ as the activity of ring neurons decreases.

The impact of LTD on the model during sleep can be understood by inspecting Figure S3B: with decreasing value of *w_EE_*, the fixed points become stable (light green) and the activity of ring neurons decreases (as shown in the fast-timescale limit). The switch between the wake and sleep phases is performed by dFB neurons that sense activity of R5 neurons^19^. Since R5 and dFB are not anatomically but funcionally connected^19^, we apply a low-pass filter to the activities of R5 neurons, which act as an input to dFB neurons and remove possible oscillations. We refer to this filtered activity as the switching signal, and it is modeled, together with dFB neurons as follows:

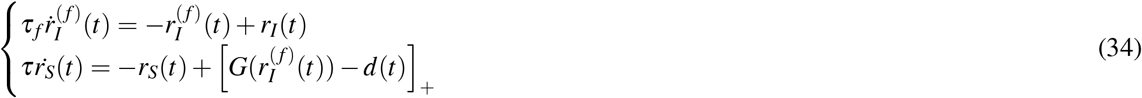

where *τ_f_* is the time constant of the low-pass filter 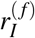 and *r_S_*(*t*) is the activity of dFB neurons with a switching behavior modeled by equation (9). The variable *d*(*t*) is a variable intended to produce sleep deprivation.

Figure S3D shows a simulation of the model combining subsequent wake (white regions), where dFB neurons are inactive, and sleep phases (grey), where dFB are active. During wakefulness, *w_EE_* and *w_IE_* undergo LTP and the activity of R5 neurons increases (light red line in second row in Figure S3D) and *r_E_* is constant. When the switching signal (dark red line in second row) crosses an upper threshold, 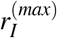, dFB neurons switch the model to sleep. During sleep, *w_EE_* undergoes LTD due to the switch in plasticity, while the activity of R5 neurons decreases. *w_IE_* also undergoes LTD (note that the plasticity rule does not change) since it follows the trend of *w_EE_* to impose the set-point *r*_0_ to the wedge neurons.

Therefore, sleep resets synaptic plasticity and activity of R5 neurons. Once the switching signal reaches a lower threshold, 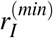, the model is switched back to the wake phase. In the third wake phase, we simulated sleep deprivation by setting *d*(*t*) = 1 (top orange region). Here, *w_EE_* crosses the bifurcation boundary, *w_EE_* > 2, and the model enters the domain of stable oscillations.

During the following sleep phase, the system needs more time to fully reset and reach the lower threshold. Such sleep rebound after sleep deprivation is an experimentally described feature of sleep homeostasis circuits^17,18^.

The time that the system spends in the sleep and wake phases is determined by the time constants of the plasticity rules, *τ_EE_* and *τ_IE_*, and the upper and lower thresholds of the switching signal, 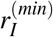 and 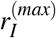. In our simulations, we set *τ_EE_* and *τ_IE_* to yield dynamics in the timescale of seconds (for ease of visualization), but larger values lead to similar behavior on longer timescales (minutes or hours, see Figure S8).

### 4.6 Ring attractor network with plasticity

We expand the two-population model to a ring attractor network. A total of *N* = 32 individual wedge neurons are modeled by 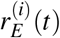. For simplicity, ring neurons are modeled as a population, *r_I_* (*t*). The model is schematically shown in Figure 2D. The dynamics of the ring attractor network are given by the following equations:

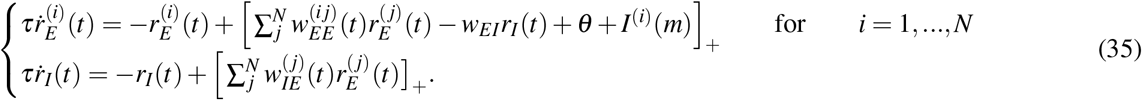

The synaptic plasticity rules are also extended from the two-population model during the wake phase:

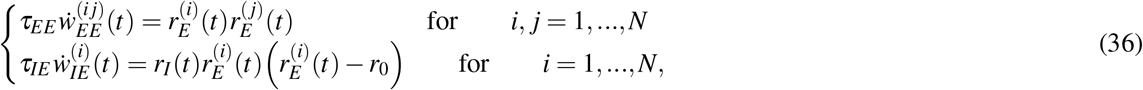

and during the sleep phase:

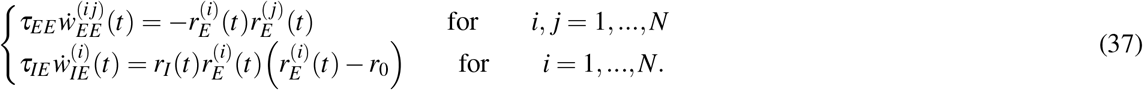

**Table S2.** Parameter values used in the ring attractor network. As in the previous model, time constants *τ, τ_f_, τ_EE_* and *τ_IE_* are measured in seconds. We consider the rest dimensionless quantities.

| $\tau$ | $\tau_f$ | $\tau_{EE}$ | $\tau_{IE}$ | $w_{EI}$ | $\theta$ | $r_0$ | $r_I^{(min)}$ | $r_I^{(max)}$ | $w_{EE}^{(max)}$ | $\sigma$ | $I_{max}$ | $I_\sigma$ |
| --- | --- | --- | --- | --- | --- | --- | --- | --- | --- | --- | --- | --- |
| 0.01 | 2 | 10000 | 10000 | 0.5 | 10 | 10 | 22 | 30 | 0.3 | 1.5 | 5 | 2 |

We initialize the synaptic weights 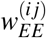 with a Gaussian function with amplitude 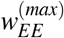 and standard deviation σ:

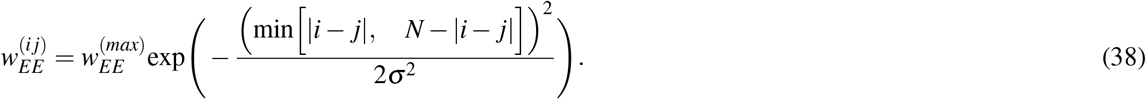

Additionally we provide a Gaussian input to the ring attractor around a given wedge neuron *m* with amplitude *I_max_* and standard deviation *I_σ_*:

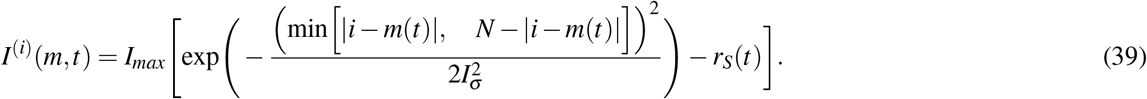

This input allows changing the position of the bump in the simulations, and can represent visual or idiothetic input to update the head direction of the animal.

We use the low-pass filtered activity of ring neurons to switch between sleep and wake phases, as in equation (5). The values of the parameters in Table S2 are used in all simulations unless stated otherwise.

Synaptic plasticity in ring and wedge neurons has been discussed in several studies^19,27^,^28^,^67^,^68^. Note that we here focus on plasticity in the connections from wedge to ring neurons, while leaving the connectivity in the opposite direction constant. This is opposite from^67^,^68^, where plasticity from ring to wedge neurons is assumed, while the other direction is left constant. This choice is motivated by the increasing activity in R5 neurons during the wake phase^19^, which could be explained by the growth of dendritic synaptic sites (pre-synaptic plasticity), for instance from wedge to R5 neurons – consistent with the data and interpretation in^19^ –, but not by the growth of axonal synaptic sites (post-synaptic plasticity).

### 4.7 Ring attractor network: bump stability analysis

To analyze to stability of the ring attractor model, we use an approach similar to the one in the fast-timescales analysis of the two-population model. First, we assume no plasticity in the recurrent connections 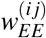 but only in 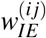,

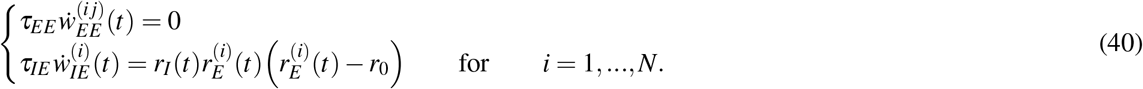

We analyze the stability and behavior of the network while gradually changing 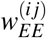. We nitialize the ring attractor network with a bump profile, where neuron number 16 has maximum activity 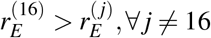. Given that only the connections 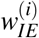 are plastic, the activity of ring neurons converges to a stable value given by

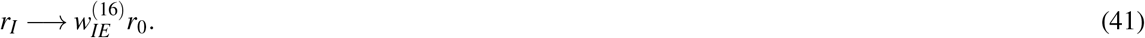

This can be understood by looking at the plasticity rule for 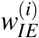 (40). First, all the synaptic weights evolve so that wedge neuron 16 approaches the activity *r*_0_. As all wedge neurons receive the same global inhibition, and wedge neuron 16 has maximum activity, the activity of the other wedge neurons is lower than *r*_0_. At this point, the weights 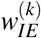 for wedge neurons *k*≠ 16 with non-zero activity, decrease over time until reaching zero. On the other hand, if a wedge neuron *q* ≠ 16 has zero activity, it does not provide any input to the ring neurons. As for wedge neuron 16, the synaptic plasticity rule changes the value of 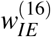 such that its activity approaches *r*_0_. An example of this behavior can be seen in Figure S5A, where we initialize the system with an input such that the peak of the bump is at wedge neuron 16. The synaptic weights 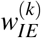 for wedge neurons *k* ≠ 16 which have non-zero activity evolve towards zero.

The value of 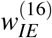 is determined by the bump profile, because 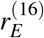 is receiving input from any wedge neuron with non-zero activity. Thus 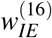 has to balance the total excitation to set the activity of the wedge neuron 16 to *r*_0_. Furthermore, the bump profile is determined by the parameters of the recurrent connectivity profile 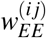, i.e. 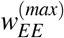 and σ (equation (38)). If we fix the amplitude 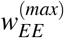 and increase the standard deviation σ, the width of the bump, that is the number of active wedge neurons, decreases. This is because more wedge neurons are providing input to wedge neuron 16 as σ increases, and the value of 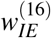 increases to set 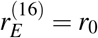, which in turn provides more inhibition through ring neurons to all the wedge neurons and lowers their activities, reducing at the same time the bump width. This behavior is seen in Figure S5A and S5B.

The bump also shows oscillatory behavior (Figure S5C) depending on the value of σ. In general, the state of the ring attractor network and its stability can be described in terms of the recurrent connectivity distribution, 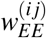, consistent with the fast-timescale limit analysis in the two-population model.

To investigate how the behavior and stability of the ring attractor network depend on the recurrent connections 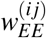, we simulated the ring attractor model for a grid of values for the parameters 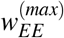 and σ:

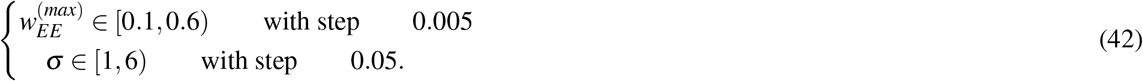

In total, we performed 10000 simulations where we initialize the bump peak in wedge neuron 16 with a predefined input of 0.5 seconds, and let the network evolve for 10 seconds. We then analyzed the stability and behavior of the system in the last second of each simulation, therefore assuming that the state of the network does not change. Figure S5 shows an example of 3 simulations with different σ values; the light orange band across all simulations highlights the region used for analysis.

In this region we computed for each simulation the following:

- **Oscillation frequency**: the frequency at which wedge neuron 16 oscillates. For this, we computed the Discrete Fourier Transform, and the resulting peak value corresponds to the oscillation frequency.
- **Mean bump FWHM**: the mean value over time of the full width at half maximum of the bump, a proxy for the width of the bump.
- **Maximum bump peak**: the maximum value of the wedge neuron 16 over time.
- **Mean bump peak**: the mean value over time of the wedge neuron 16. If the bump does not oscillate, this value is equivalent to the maximum bump peak value.
- **Mean ring neuron activity**: the mean of ring neuron activity over time.

These measures are displayed in Figure S6B,S6C,S6D, S6E and S6F, as a function of the recurrent connectivity parameters 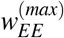 and σ. Figure S6B shows how the network starts oscillating with increasing 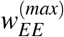 and σ. On the other hand, the FWHM in Figure S6B and S6C shows how at low 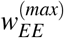 and σ values, the bump disappears and all wedge neurons have constant activity at *r*_0_. As the parameter values increase, a bump of activity appears and the FWHM decreases, as observed in Figure3C and S5A and B. Finally, the activity of ring neurons increases as 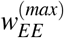 and σ increase.

We further characterise the behavior and stability of the bump in the network with the total excitatory connectivity to the wedge neuron with maximal activity, i.e. wedge neuron 16, 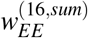:

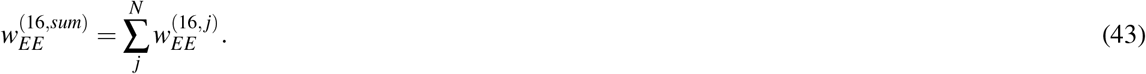

Figure S6A highlights the isolines where 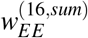 is constant for different values of 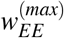 and σ. Note how the constant values of 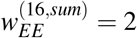 and 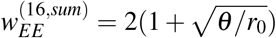 coincide with the boundaries of the different dynamic regimes. Therefore, the quantity 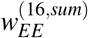 has in the ring attractor model a similar role as does the recurrent connection *w_EE_* in the two-population model. For 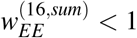, the bump disappears, similar to the two-population network model when *w_EE_* < 1. However, unlike in the two-population model, which is unstable for 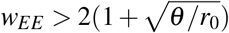, in the ring attractor network, for 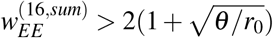 the wedge neurons are strongly inhibited by high activity in ring neurons.

From the above analysis we extracted regions of stability that are shown in Figure 3B. Constant lines of 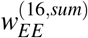 are computed as follows: in the continuous limit, i.e. *N* → ∞, the total excitatory connectivity is given by the following integral:

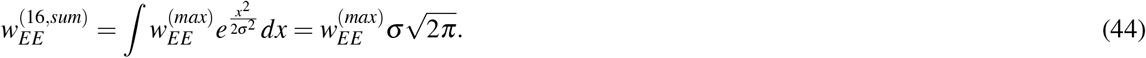

Therefore, the isolines of constant 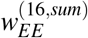 are given by 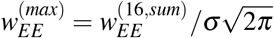. For the discrete case, we empirically found 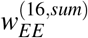 to be well approximated by:

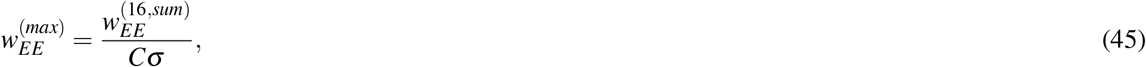

for any constant line 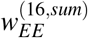, where *C* ≈ 2.697.

### 4.8 Ring attractor network: autonomous bump path analysis

To simulate how the bump in the ring attractor changes position to update the head direction during the wake phase, we use a simple clockwise or counter-clockwise rotating input with frequency *f* defined by:

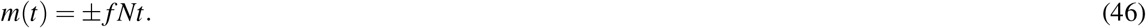

*m*(*t*) is the wedge neuron where the ring attractor receives the Gaussian input *I^(i)^(m(t))* (equation (39)), and it is a cyclic variable, so that:

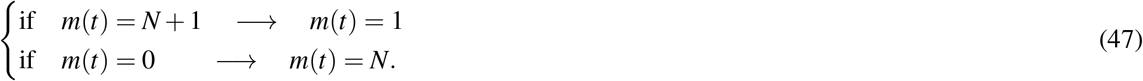

During the sleep phase, the bump in the ring attractor shows autonomous dynamics (Figure 5A). To investigate the relationship between the path of the bump during sleep and in the preceding wake phase, we use a sinusoidal input during the wake phase, defined by the amplitude *A*, frequency *f*, and the center *C*:

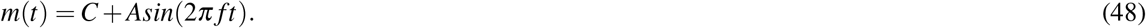

An example of this input with different frequencies *f* and centers *C* is shown in Figure 6A. We can obtain the position of the bump during wake and sleep phases as:

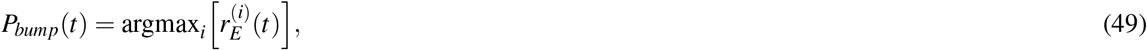

so that the position of the bump corresponds to the wedge neuron with maximum activity. We can now compute the distribution of times that the bump is localized around each wedge neuron *i* during sleep and wake phases, respectively, as:

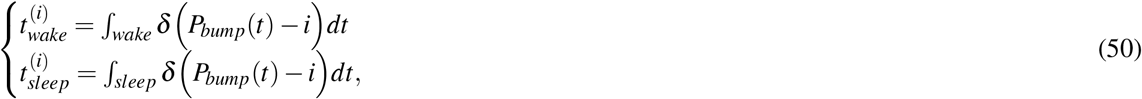

where δ(·) is the Dirac delta function and integrals extend over the wake and sleep phases. Figure 6B shows these distributions normalized for the three wake phases in Figure 6A and their following sleep phases. The distributions are very similar, meaning that during the sleep phase, the bump revisits the same wedge neurons that were active during the wake phase.

We further asked how the autonomous bump path changes during sleep with respect to the amplitude of the sine-shaped input, A, and frequency, *f*, during sleep. We first fixed the frequency of the input at *f* = 1*Hz* and the center at *C* = 16 while varying the amplitude in the range of [0,15] with an increment of 1. This resulted in 15 simulations where we computed the standard deviation of the bump path during sleep and wake:

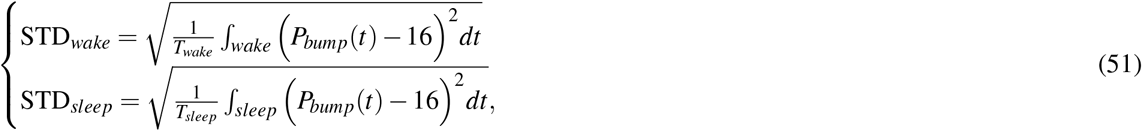

where *T_wake_* and *T_sleep_* are the duration of wake and sleep phases, respectively. Figure 6C shows the standard deviation in both phases with respect to the amplitude *A*. Note the similarity between both phases.

Secondly, we fixed the value of the amplitude at *A* = 8 and the center at *C* = 16 while varying the frequency, *f*, in the range [0.1,1.5] at increments of 0.1*Hz*, resulting in 15 simulations. We quantify the number of cycles during both the sleep and wake phase. During the wake phase, the number of cycles is proportional to the input frequency *f*. Therefore a linear relationship between the number of cycles during sleep and the frequency of the input would give correlation between the frequency and the autonomously rotating bump path. However, Figure 6C, bottom, shows how the number of cycles during sleep does not change as the input frequency increases. An example of this can be seen in Figure 6A where we increase the frequency in consecutive wake phases and the path of the bump during sleep does not increase the rotation frequency.

### 4.9 Ring attractor network: bump drift during wake phase

In the simulations and analyses above, we provided input during the wake phase and the ring attractor network closely followed the input with a bump of activity. However, a ring attractor network should be able to sustain the bump of activity in the absence of input. It is known that small changes in the synaptic connections of wedge neurons 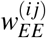 can cause drifts of the bump in the absence of input^43-45^.

To test for drift during the wake phase, we used a flashing rotating input that turns on and off. The input around a wedge neuron *m* is on for 0.2 sec (equation (39)), and then is turned off for 0.3 sec. Therefore, for *N* neurons the rotating input frequency is 1/(0.5*N*). Figure S8A shows such a simulation with three wake and sleep phases and S8B, C and D show zoom-ins around different times. Note how the bump drifts from the provided visual input, due to the synaptic changes in 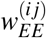.

This drift depends on the plasticity time constants. For instance, Figure S9 shows a simulation with 100 times larger time constants, *τ_EE_, τ_IE_* = 10^6^. The duration of wake and sleep phases are now in the order of hours, compared to the simulation in Figure S9. Note in the zoom-ins of Figures S9B, C and D that the bump of activity is sustained during the off time of the visual input without drifting. Since there are many more rotations during the wake phase and the synaptic changes in 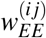 are very small in each rotation, the weights increase all together very homogeneously.

**Figure S1.**
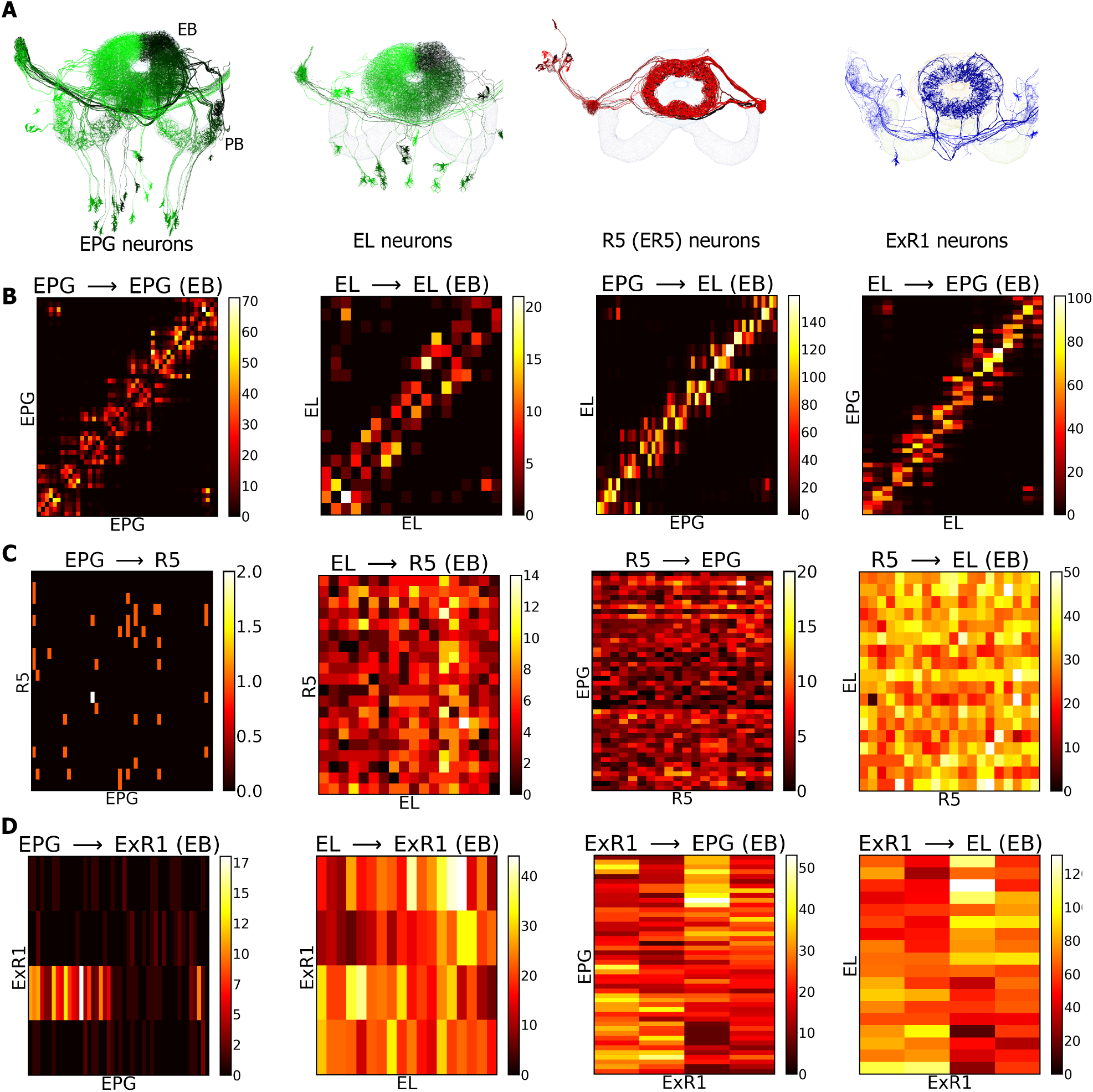
Connectivity between the navigation-related populations (EPG and EL) and sleep-related populations (R5 and ExR1). **A** On the left, neural projections of EPG and EL, referred as wedge neurons (green). On the right, neural projections of R5 (red) and ExR1 (blue). **B** Recurrent connectivity between wedge neurons. The matrix in each figure represents the number of synaptic sites between presynaptic neurons (horizontal axis) and postsynaptic neurons (vertical axis)^14^. **C** Connectivity between wedge neurons and R5 neurons in both directions. **D** Connectivity between wedge neurons and ExR1 neurons in both directions. Data and neurons are reproduced from^14^.

**Figure S2.**
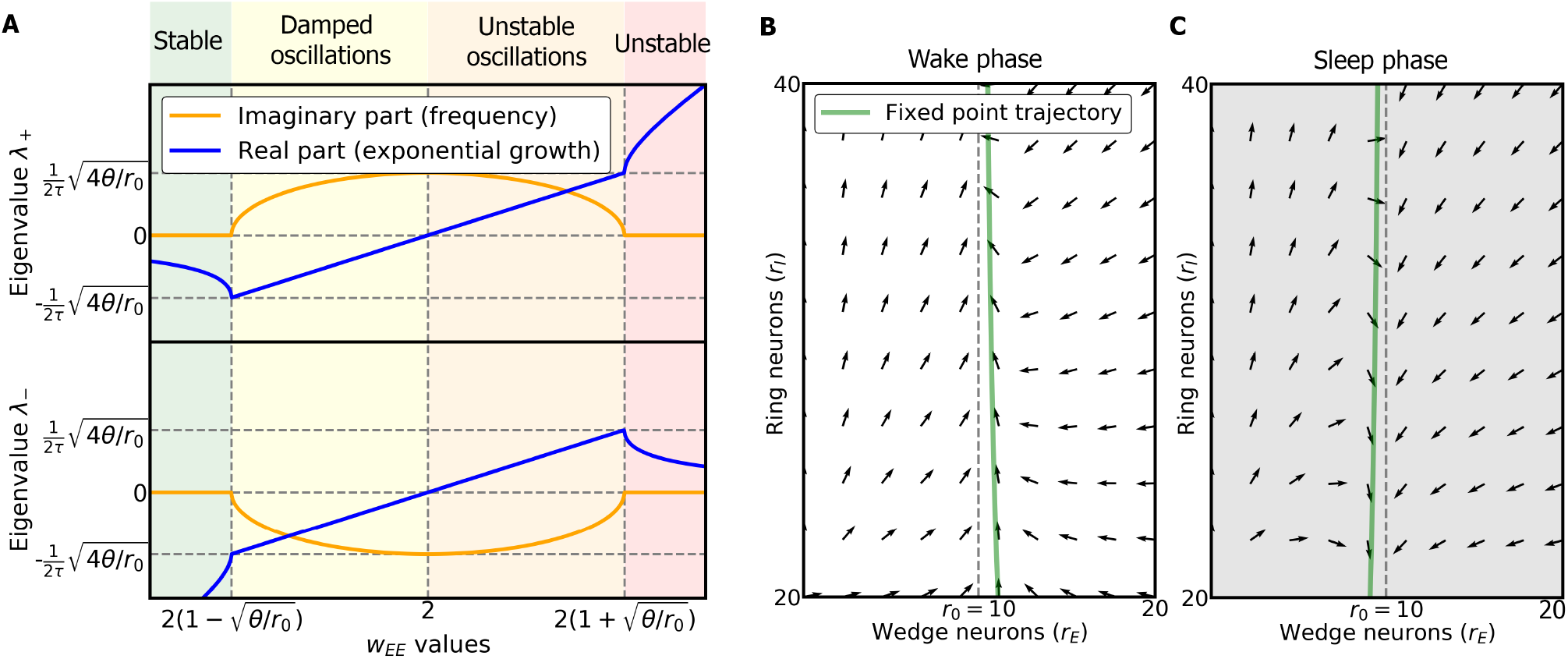
Analysis of the two population model in the fast and slow-timescale limits. **A** Eigenvalues in the two-population model in the fast-timescales limit. The real (in blue) and imaginary (in orange) parts of the eigenvalues are plotted as a function of *w_EE_*. **B** Vector field of wedge and ring neurons dependent on synaptic plasticity during the wake phase in the slow-timescale limit. The green line represents the set point trajectory of wedge neurons. **C** Vector field of wedge and ring neurons due to plasticity in the sleep phase in the slow-timescale limit. The green line is the trajectory of the set point in wedge neurons.

**Figure S3.**
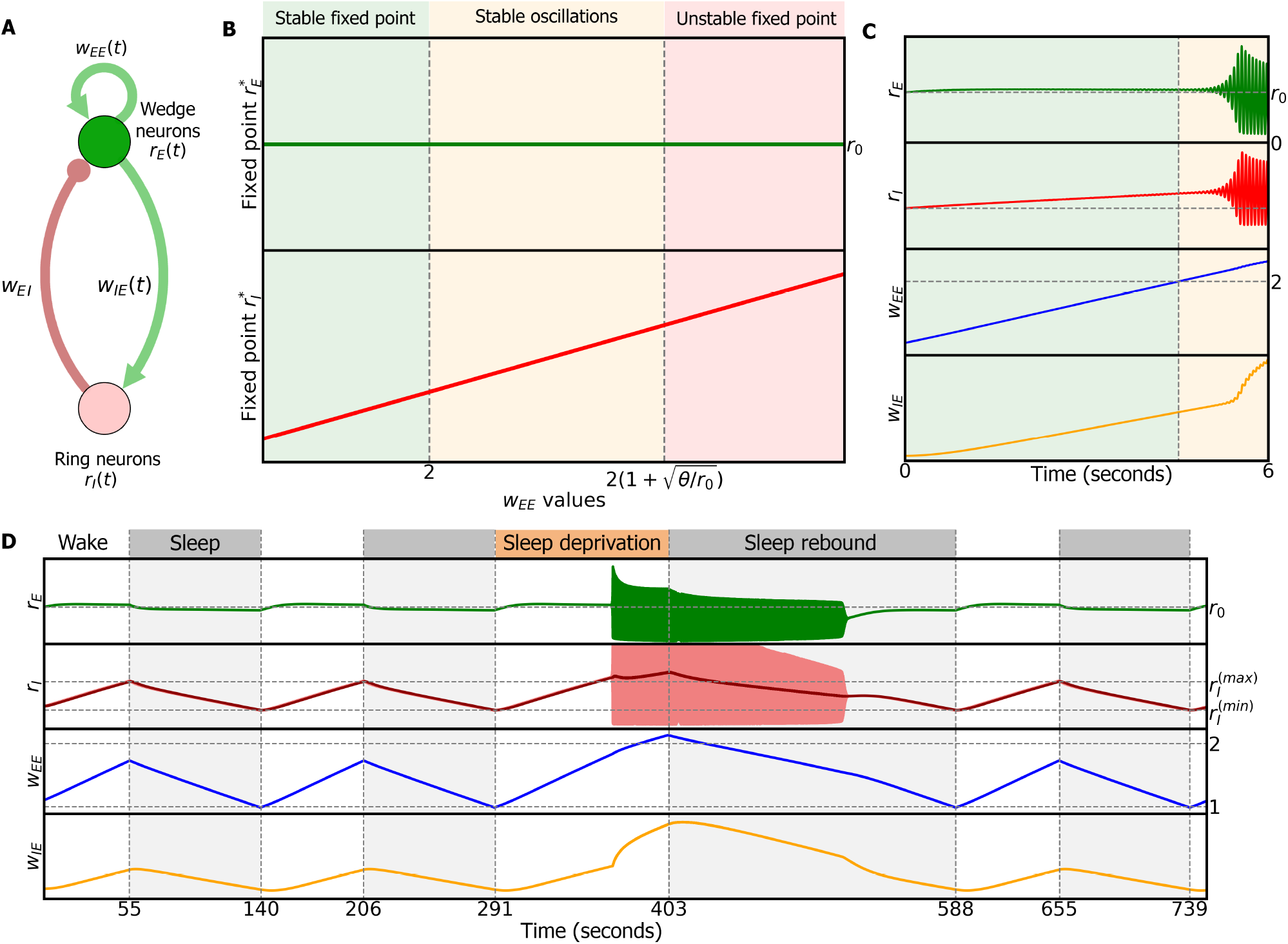
Two-population model. **A** Model describing the dynamics of the activities of excitatory wedge neurons and inhibitory ring neurons, *r_E_* and *r_I_* respectively, and the plasticity of the synaptic weights within the wedge population *w_EE_* and from the wedge population to the ring population *w_IE_*. Green connections are excitatory, red connections inhibitory. **B** Stability conditions with respect to *w_EE_* (fast-timescale limit, see Methods). **C** Dynamics during wake phase. As *w_EE_* grows, the system undergoes a Hopf bifurcation (a critical point where the system starts to oscillate) and both populations start to oscillate around a fixed point. **D** Dynamics with alternating sleep and wake phases. Wake phase produces LTP in *w_EE_* and *w_IE_* and increases the activity of ring neurons. Sleep produces LTD and reduces the activity of ring neurons. Extending the wake period produces sleep deprivation and results in stable oscillations. The subsequently required sleep period for resetting is longer (sleep rebound).

**Figure S4.**
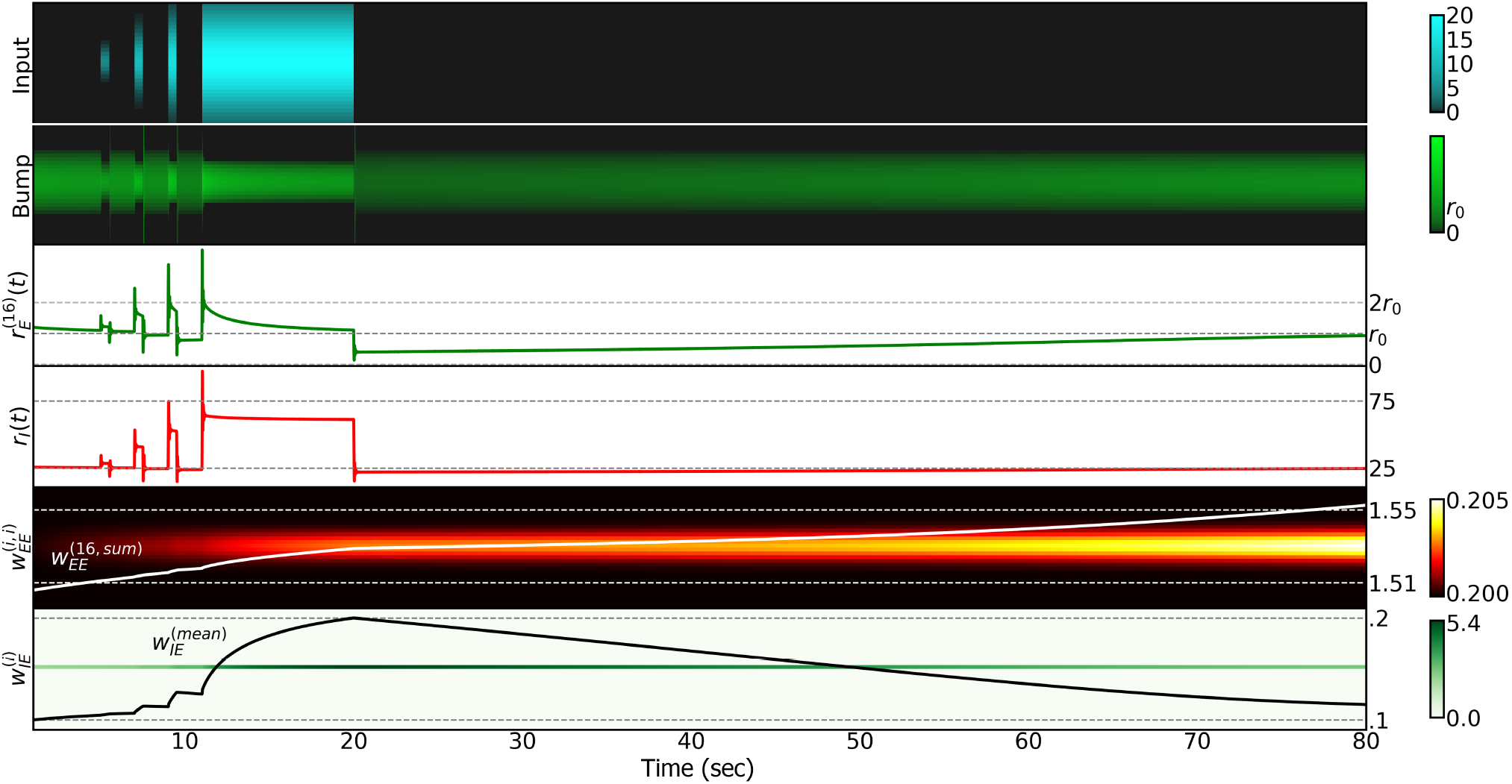
Bump profile in the ring attractor network in response to inputs of different amplitudes *I_max_* and standard deviations *I_σ_*. When input (first row) is provided to the ring attractor the plasticity rule for 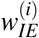 brings the activity of the wedge neuron 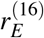 (where the bump peak is located, second row) back to *r*_0_ (third row). The bump is not constrained to have constant activity (third row), but always relaxes towards *r*_0_ over time in the absence of changing input. For ease of visualization, we used a slower time constant for the plasticity of 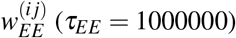, thus avoiding oscillations in the bump throughout the simulation.

**Figure S5.**
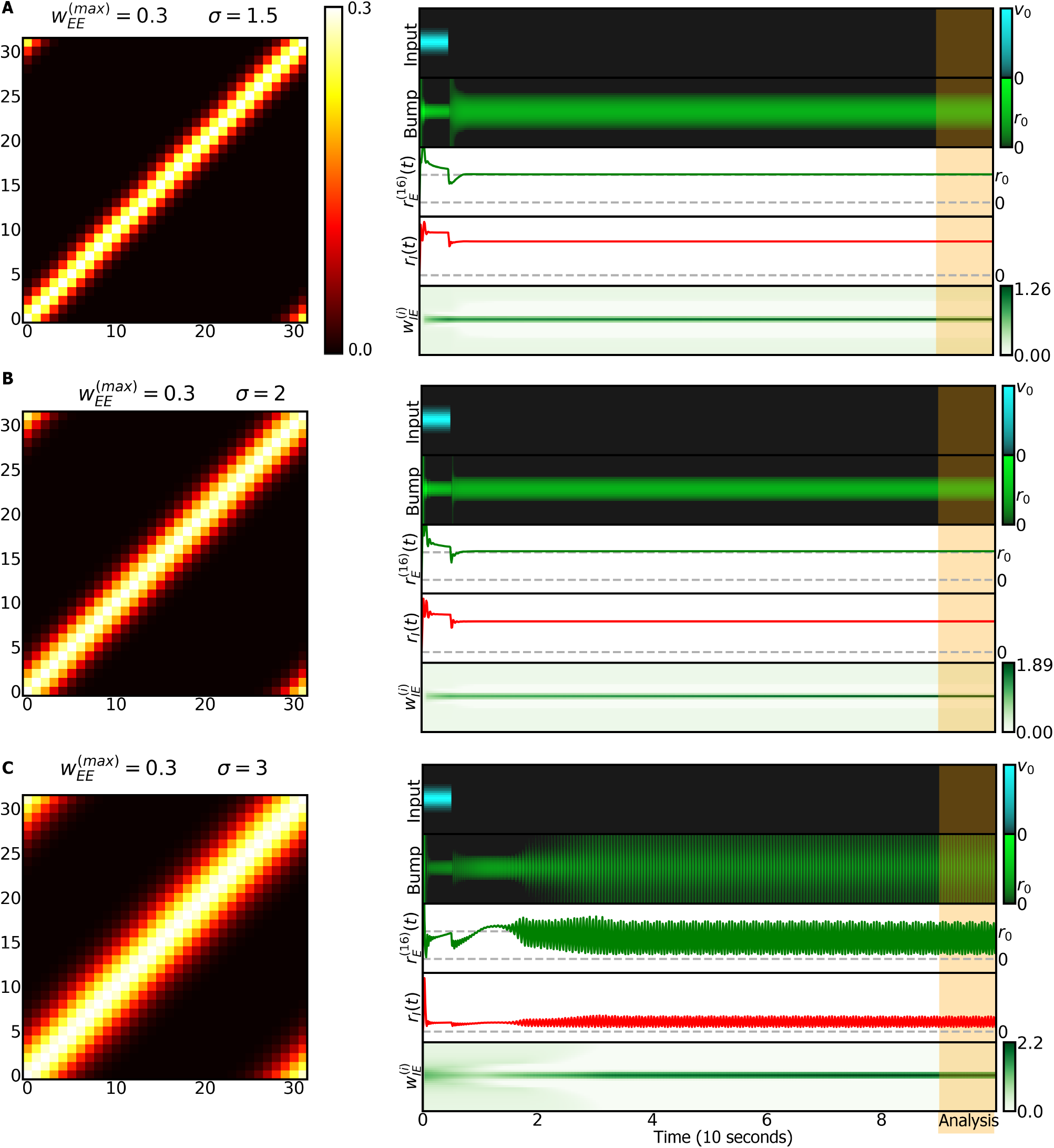
Ring attractor network with no plasticity in 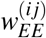 and different initial distributions of weights. **A** Left: initial 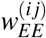 values; right: 10 seconds of simulated dynamics. First row: input positioning the bump around wedge neuron 16. Second row: bump profile over time. Third row: activity of wedge neuron 16 evolving towards *r*_0_ due to plasticity in 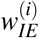. Fourth row: activity of ring neurons. Fifth row: the synaptic weights 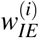. The orange band at the end of the simulations represents the period in which the stability of the bump is analyzed. **B** Same as A but with larger σ. **C** Same as A, B but again increasing σ. In this simulation, the bump shows stable oscillations.

**Figure S6.**
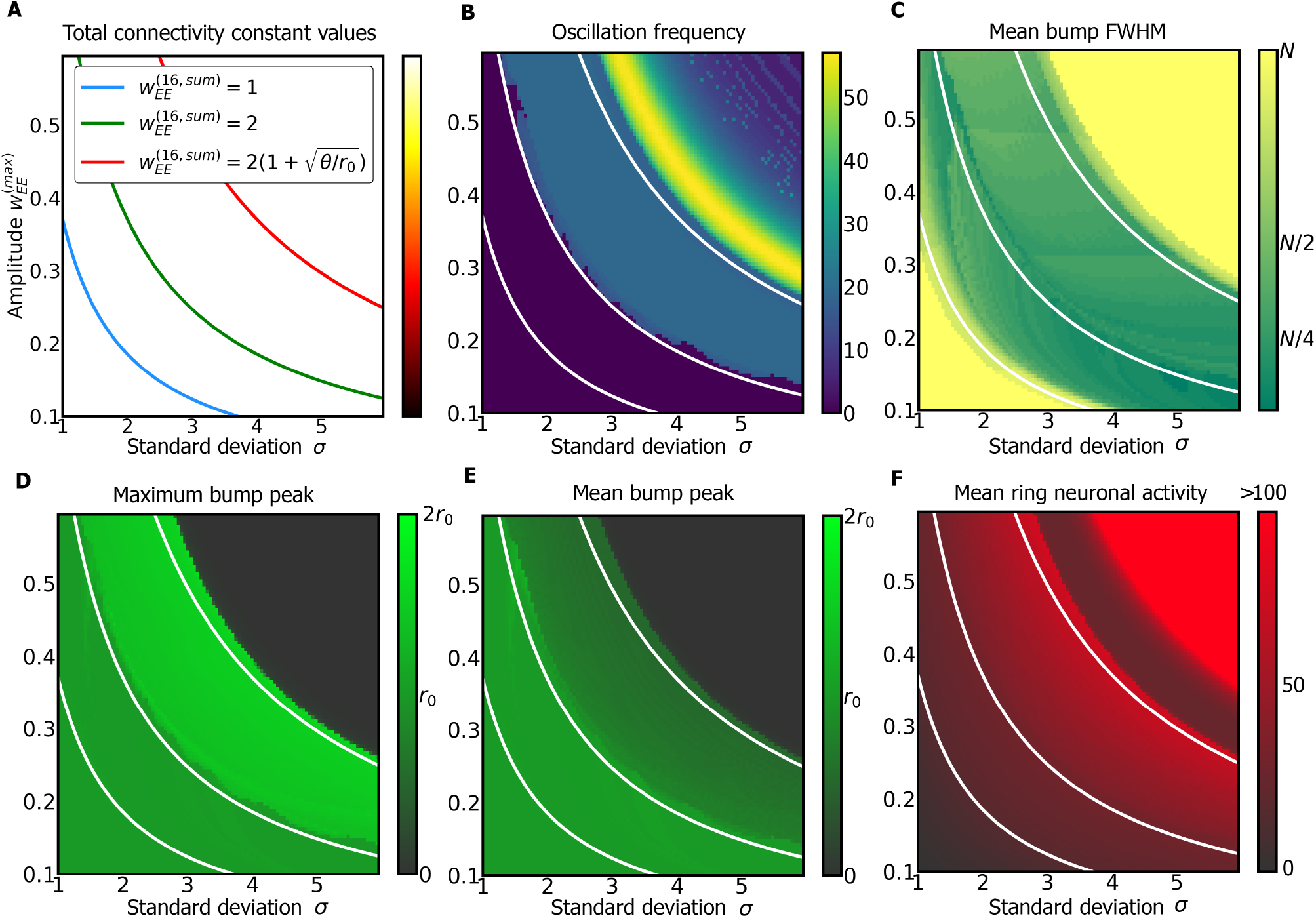
Stability analysis of the bump in the ring attractor network. All subfigures are plots of different measures as a function of the grid of values of σ and 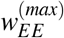 that define the fixed synaptic weights 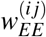 in the bump stability analysis (Methods (4.7)). All variables in B, C, D, E, and F are calculated in a time slot of 1 sec at the end of each simulation (orange area in Figure S3). **A** Isolines of 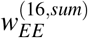. These isolines are overlaid in white in the other subfigures and represent approximate boundaries of stability. **B** Frequency of the oscillations in the bump across the analysis time slot. Above the isoline 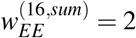, the bump oscillates. **C** Mean FWHM over the analysis time slot. Below the isoline 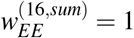 and above the isoline 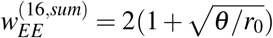 there is no bump because the FWHM is equal to the number of neurons, *N*. **D** Maximum peak of the bump in the analysis time slot (maximum activity of wedge neuron 16, 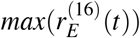). Below the isoline 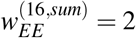, the maximum is *r*_0_ as forced by the plasticity rule in 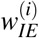. Between the isolines 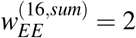 and 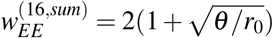, the bump oscillates and the amplitude is given by the maximum, in this case 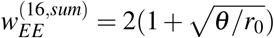, there is close-to-zero activity in wedge neurons, since the maximum is near zero. **E** Mean of the bump peak over the analysis slot time (mean activity of the wedge neuron 16, 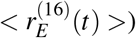. In case of no oscillations, this value should be equivalent to the maximum bump peak in panel D. During oscillations, the value closely represents the center of oscillations. Note that between isolines 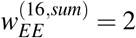 and 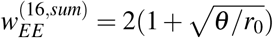 this value is lower than *r*_0_. **F** Mean activity over the analysis time slot of ring neurons, < *r_I_*(*t*) >. Below the isoline 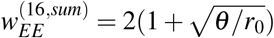 the activity of ring neurons increases. Above the isoline, the activity of ring neurons first decreases and then rapidly increases. We clipped the values of the mean activity above 100 to facilitate visualization, but the increment of activity in this area reached values over 1000.

**Figure S7.**
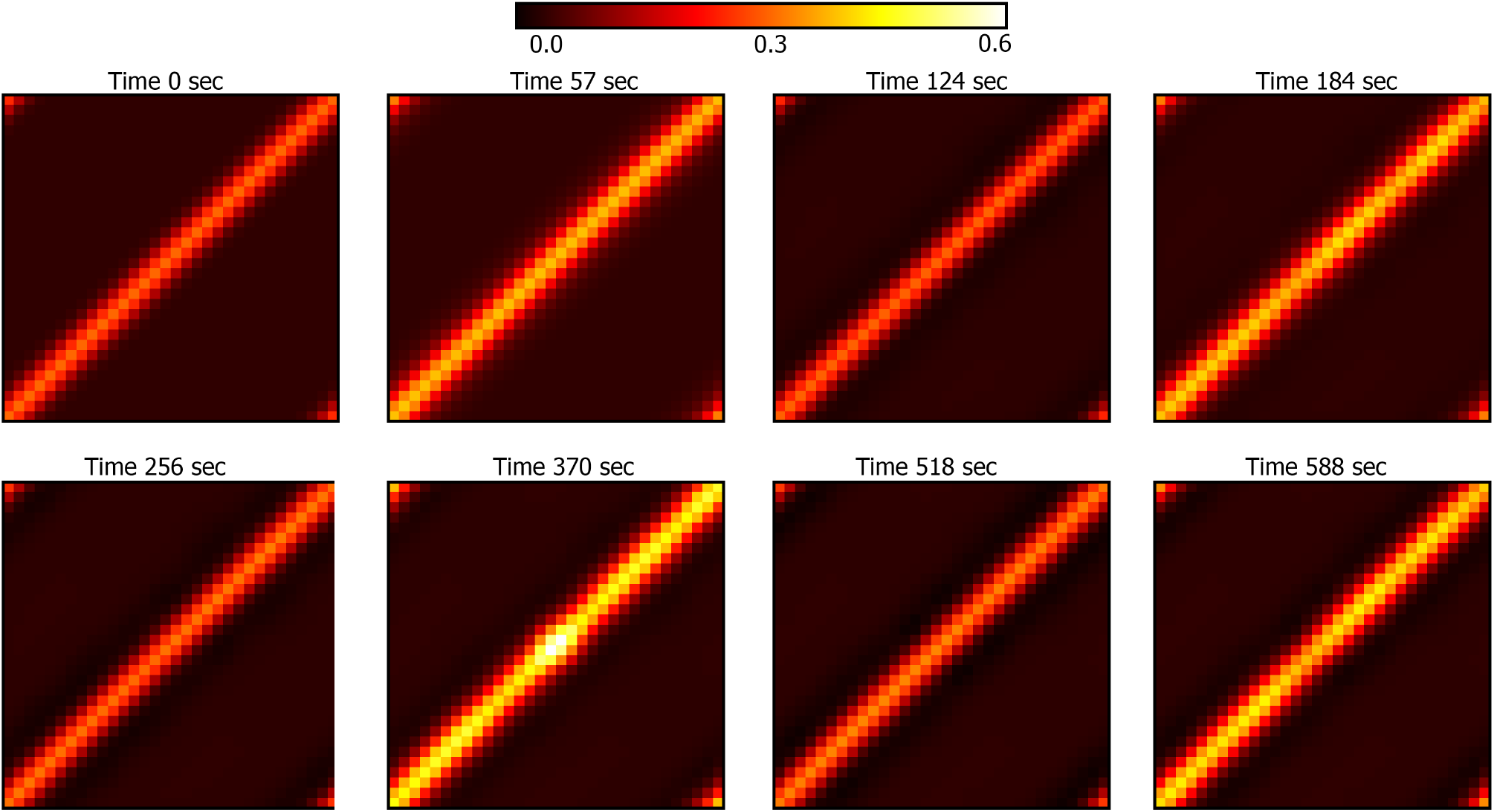
Values of the synaptic weights 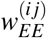 in the simulation of Figure 3. From left to right and from top to bottom: each plot shows the values each time there is a switch between wake and sleep phase and *vice versa*. Top, left: initial values of the weights set by the initial values of 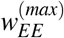 and σ in Table S2. Top, center left: weights at the end of the first wake phase. Top, center right: after the subsequent sleep phase. Top, right: after the second wake phase. Bottom, left: weights after the second sleep phase. Bottom, center left: after sleep deprivation. Bottom, center right: after sleep rebound. Bottom, right: weights after the third wake phase. Note how the weights after each wake phase are increased (specially after sleep deprivation in the sixth plot) and how the sleep phase resets the weights to close to the initial conditions.

**Figure S8.**
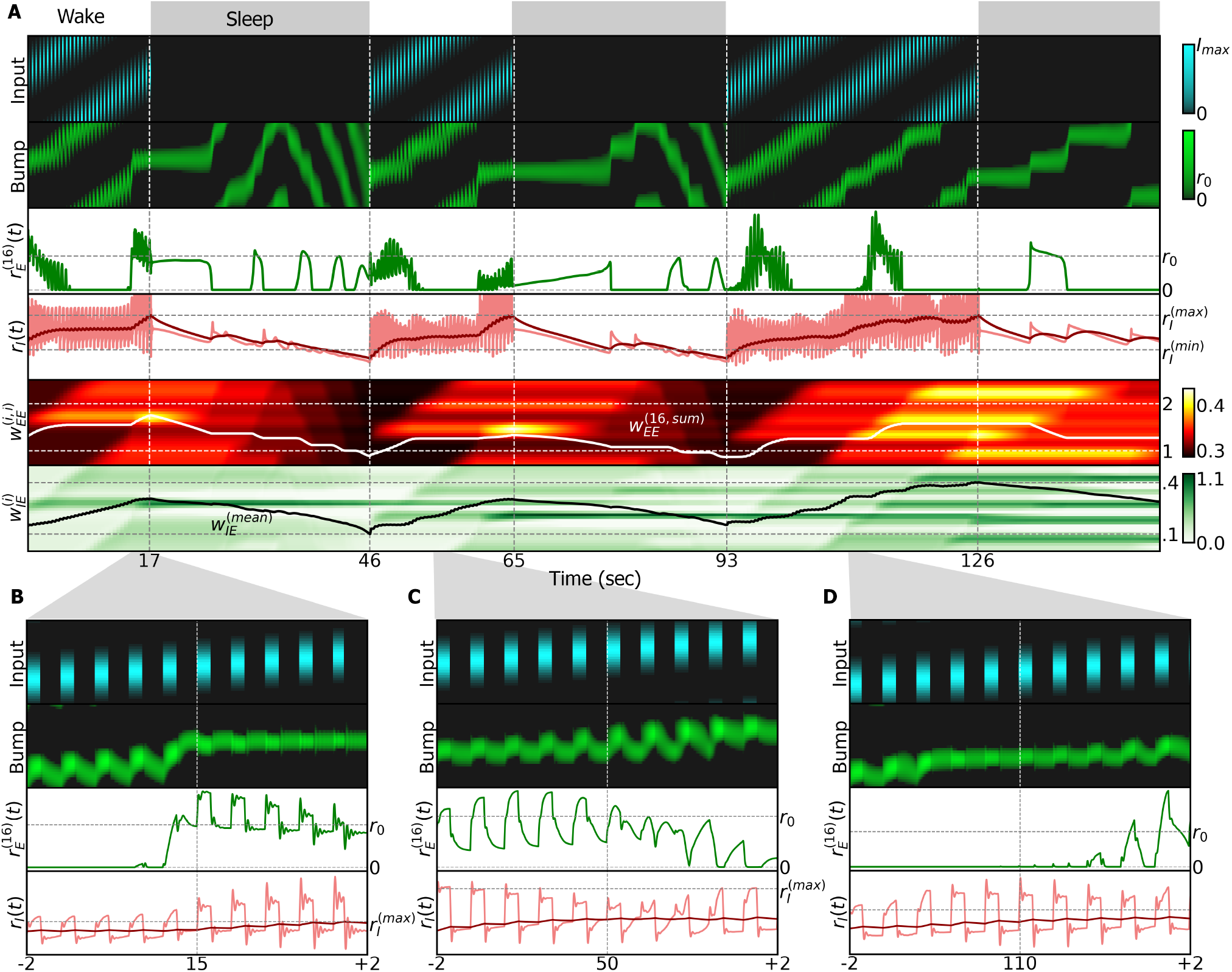
Simulation of the ring attractor network with flashing input during the wake phase (details in Methods(4.9)). **A** Simulation with three wake phases and three sleep phases. A flashing input that turns on and off is provided during the wake phase. During the off period, the ring attractor sustains a bump of activity that drifts due to changes in the synaptic weights 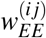. **B** Zoom-in around 15 seconds in the first wake phase of the simulation. **C** Zoom-in around 50 seconds in the second wake phase. **D** Zoom-in around 110 seconds in the third wake phase.

**Figure S9.**
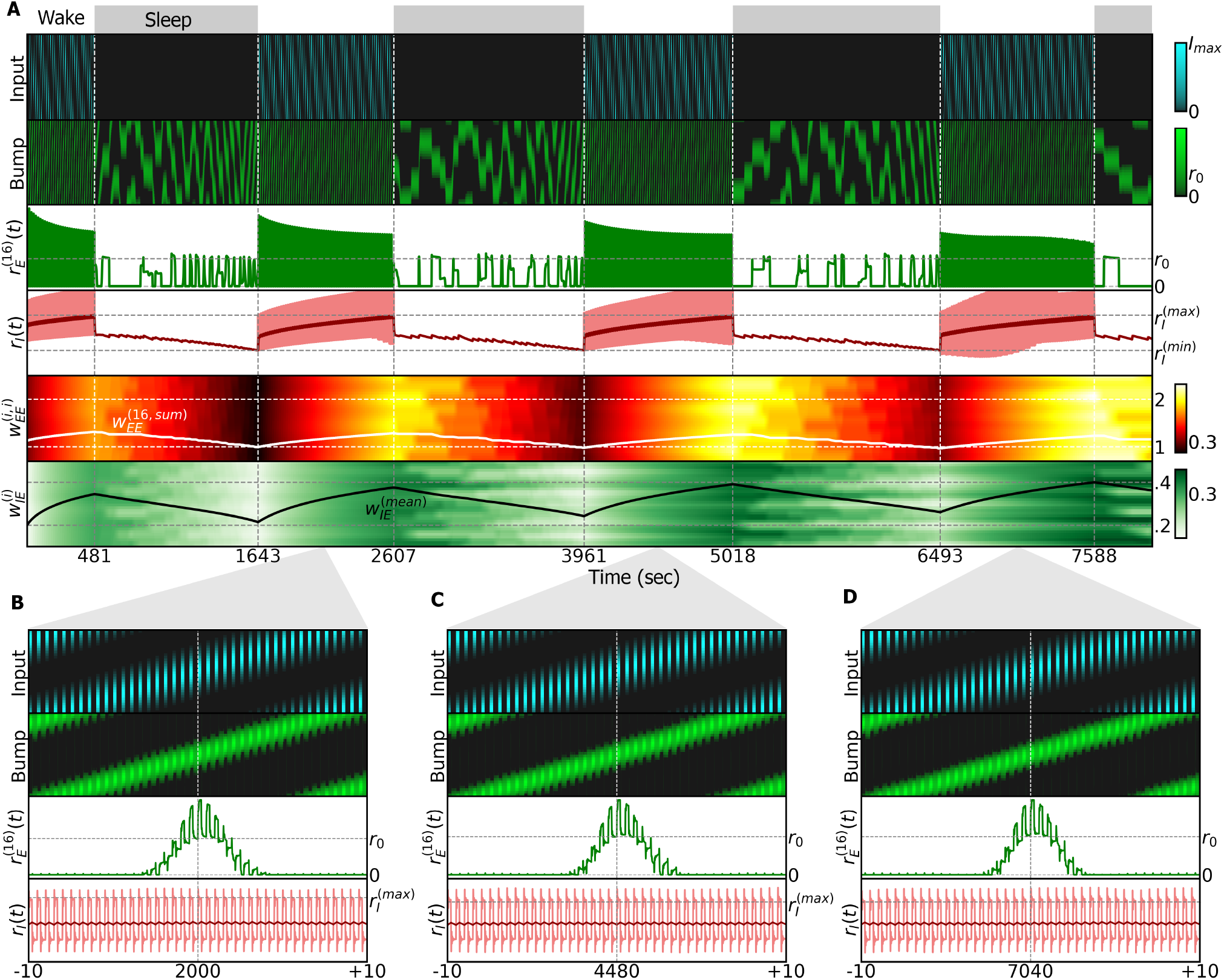
Simulation of the ring attractor network with flashing input during the wake phase with slow plasticity time constants (details in Methods(4.9)). The time constants in this simulation are 100 times larger than in the previous ring attractor simulations. **A** Full simulation with three wake phases and three sleep phases. We used the same input rotation frequency as in Figure S7. As in Figure S7, we provide a flashing input that turns on and off during the wake phase. **B** Zoom-in around 2000 seconds in the first wake phase of the simulation. **C** Zoom-in around 4480 seconds in the second wake phase. **D** Zoom-in around 7040 seconds in the third wake phase. Note how the duration of sleep and wake phases are now in the order of hours due to the increased time constants in the plasticity rules. Note also how the bump of activity is sustained in place without drifting after the input switches off.

